# Rapid discovery of synthetic DNA sequences to rewrite endogenous T cell circuits

**DOI:** 10.1101/604561

**Authors:** Theodore L. Roth, P. Jonathan Li, Jasper F. Nies, Ruby Yu, Michelle L.T. Nguyen, Youjin Lee, Ryan Apathy, Anna Truong, Joseph Hiatt, David Wu, David N. Nguyen, Daniel Goodman, Jeffrey A. Bluestone, Kole Roybal, Eric Shifrut, Alexander Marson

## Abstract

Genetically-engineered immune cell therapies have been in development for decades^1–3^ and recently have proven effective to treat some types of cancer^4^. CRISPR-based genome editing methods, enabling more flexible and targeted sequence integrations than viral transduction, have the potential to extend the clinical utility of cell therapies^5,6^. Realization of this potential depends on improved knowledge of how coding and non-coding sites throughout the genome can be modified efficiently and on improved methods to discover novel synthetic DNA sequences that can be introduced at targeted sites to enhance critical immune cell functions. Here, we developed improved guidelines for non-viral genome targeting in human T cells and a pooled discovery platform to identify synthetic genome modifications that enhance therapeutically-relevant cell functions. We demonstrated the breadth of targetable genomic loci by performing large knock-ins at 91 different genomic sites in primary human T cells, and established the power of flexible genome targeting by generating cells with Genetically Engineered Endogenous Proteins (GEEPs) that seamlessly integrate synthetic and endogenous genetic elements to alter signaling input, output, or regulatory control of genes encoding key immune receptors. Motivated by success in introducing synthetic circuits into endogenous sites, we then developed a platform to facilitate discovery of novel multi-gene sequences that reprogram both T cell specificity and function. We knocked in barcoded pools of large DNA sequences encoding polycistronic gene programs. High-throughput pooled screening of targeted knock-ins to the endogenous T cell receptor (TCR) locus revealed a transcriptional regulator and novel protein chimeras that combined with a new TCR specificity to enhance T cell responses in the presence of suppressive conditions *in vitro* and *in vivo*. Overall, these pre-clinical studies provide flexible tools to discover complex synthetic gene programs that can be written into targeted genome sites to generate more effective therapeutic cells.

## RESULTS

We recently developed an efficient non-viral genome targeting method to modify endogenous genes in primary human T cells^6,7^. Unlike non-targeted gene transduction with viral vectors, CRISPR-based knock-in requires a combination of gRNA and homology-directed repair (HDR) template sequences to target a transgenic DNA sequence to a specified genomic site. Gene targeting efficiencies vary significantly from locus to locus^6^. To determine the spectrum of endogenous genomic loci amenable to non-viral genome targeting, we knocked-in large (∼800bp) reporter constructs, encoding GFP or a truncated nerve growth factor receptor (tNGFR), into 91 unique genomic sites in cells from six healthy human blood donors (**Fig. 1a)**. The screen was performed in both CD4+ and CD8+ T cells with protein knock-in efficiencies assessed in both resting and re-stimulated cells (**Extended Data Fig. 1, 2**). Targeting of diverse genes encoding T cell receptor (TCR) complex members, immune surface receptors, checkpoint receptors, transcription factors, and many cytoskeletal and housekeeping genes revealed a wide range of observed knock-in efficiencies (**Fig. 1a** and **Extended Data Fig. 1**). Nonetheless, 88% of the loci targeted had observed knock-in efficiencies >5% with at least one gRNA (median efficiency of 21.3%). We found that RNA expression of the endogenous target gene and DNA accessibility at the gRNA cut site were both correlated with observed knock-in efficiency (**Extended Data Fig. 3**). A multivariate linear regression analysis based on gRNA sequence, RNA expression of the target gene, and chromatin accessibility helped explain variation in knock-in efficiency at diverse genomic sites (**Fig. 1b** and **Extended Data Figs. 3, 4**).

**Figure 1:**
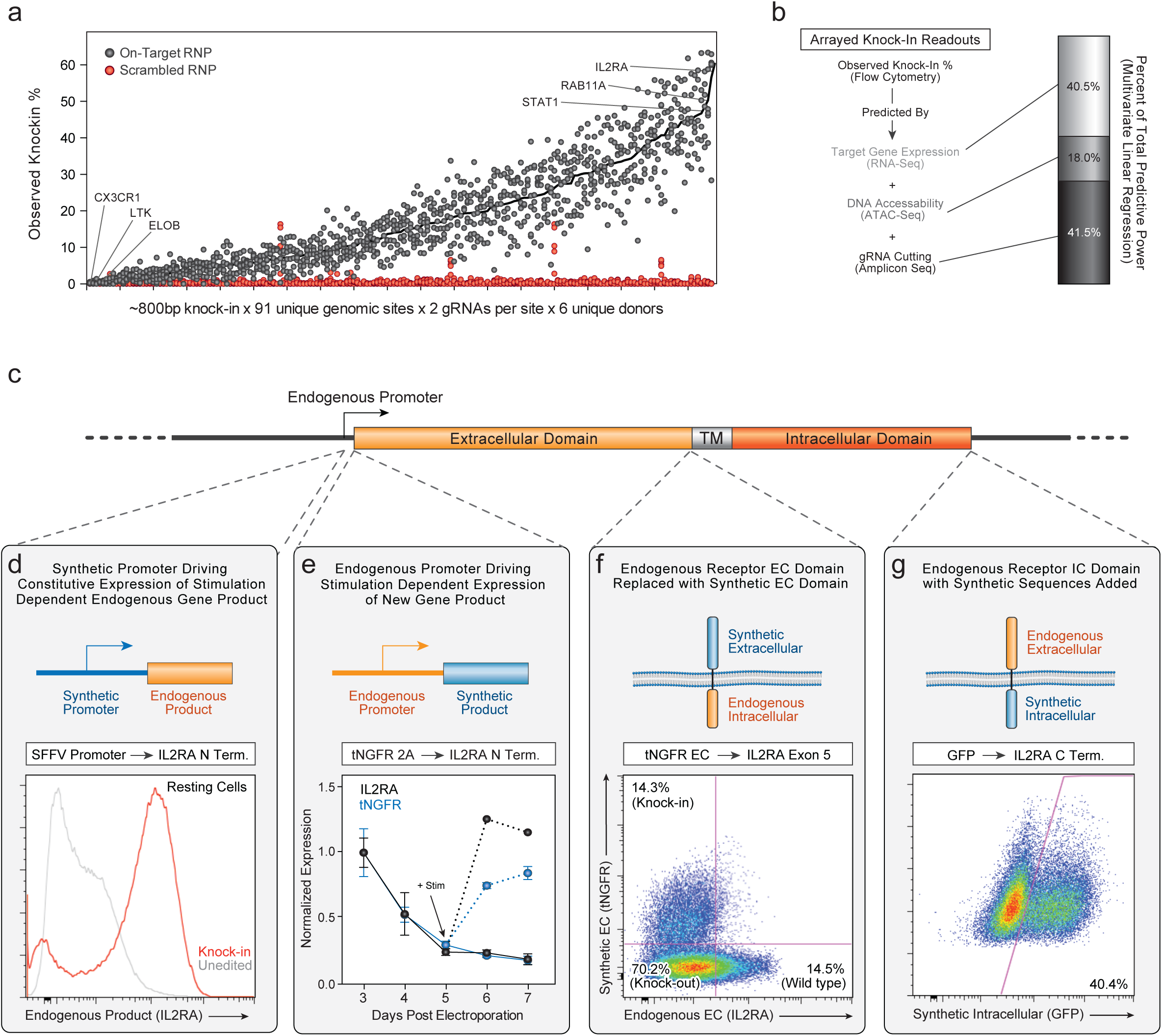
Arrayed knock-ins across endogenous loci reveal guidelines for efficient non-viral gene targeting in primary human T cells. **a**, Observed knock-in percentages for knock-in of a large HDR templates (either GFP or tNGFR, ∼800 bps) across 91 unique genomic target sites, testing two gRNAs per site. A wide range of knock-in efficiencies were observed, from below detectable levels at genes such *CX3CR1* and *ELOB* to averages across donors of ∼50% at *IL2RA, RAB11A*, and *STAT1* (overall median knock-in rate was 21.3%). Across tested sites, observed knock-in was much higher with on-target RNPs (black) than with a scrambled RNP (red) that does not target any human genomic sequence. **b**, Observed knock-in percentages were analyzed by flow cytometry. RNA-seq, ATAC-seq, and amplicon sequencing were performed for each blood donor (**Extended Data Fig. 2**). A multivariate linear regression model’s weighting of individual parameters revealed large independent contributions to observed knock-in % from RNA expression levels of the target gene, DNA accessibility at the gRNA target site, and gRNA observed insertion/deletion mutation (indel) efficiency. An ideal target genomic locus for non-viral CRISPR targeting in primary T cells is highly expressed, accessible at the time of electroporation, and contains a target sequence for a gRNA that cuts efficiently **c**, Schematic description of strategies for engineering a gene locus to generate Genetically Engineered Endogenous Protein (GEEP) primary T cells. An endogenous gene encoding a critical immune receptor (IL2RA) can be modified to reprogram: 1) the gene regulation, 2) the gene product, 3) the extracellular domain, or 4) the intracellular components. **d**, To reprogram endogenous gene regulation of IL2RA, we targeted an SFFV promoter to the gene’s 5’ non-coding region. When cultured without restimulation for 11 days following electroporation, CD4+ T cells with the SFFV promoter knock-in (Knock-in) showed sustained, stimulation-independent expression of IL2RA assessed by flow cytometry. In contrast, control CD4+ T cells without SFFV promoter knock-in (Unedited) showed low IL2RA surface expression, consistent with expected expression dynamics of endogenous IL2RA in resting cells. **e**, We demonstrated placement of a synthetic/exogenous product under endogenous gene regulatory circuits by targeting a tNGFR-2A construct to the start codon of *IL2RA* for co-expression of tNGFR and IL2RA. Starting on Day 3 post electroporation, we analyzed the expression dynamics of both tNGFR and IL2RA in edited CD4+ T cells by flow cytometry. Surface expression of both IL2RA and tNGFR decreased over time in the absence of restimulation. Upon CD3/CD28 restimulation, tNGFR surface expression was induced along with IL2RA in edited CD4+ T cells. **f**, We demonstrated replacement of an extracellular domain by targeting the extracellular domain of tNGFR to Exon 5 of the endogenous *IL2RA* gene. Upon restimulation, edited CD4+ T Cells stained for tNGFR but not IL2RA (Knock-in). **g**, The intracellular domain of endogenous *IL2RA* could be modified with addition of a c-terminus with a GFP fusion. Representative plots and graphs from n=2 healthy donors (**d-g**).

**Figure 2:**
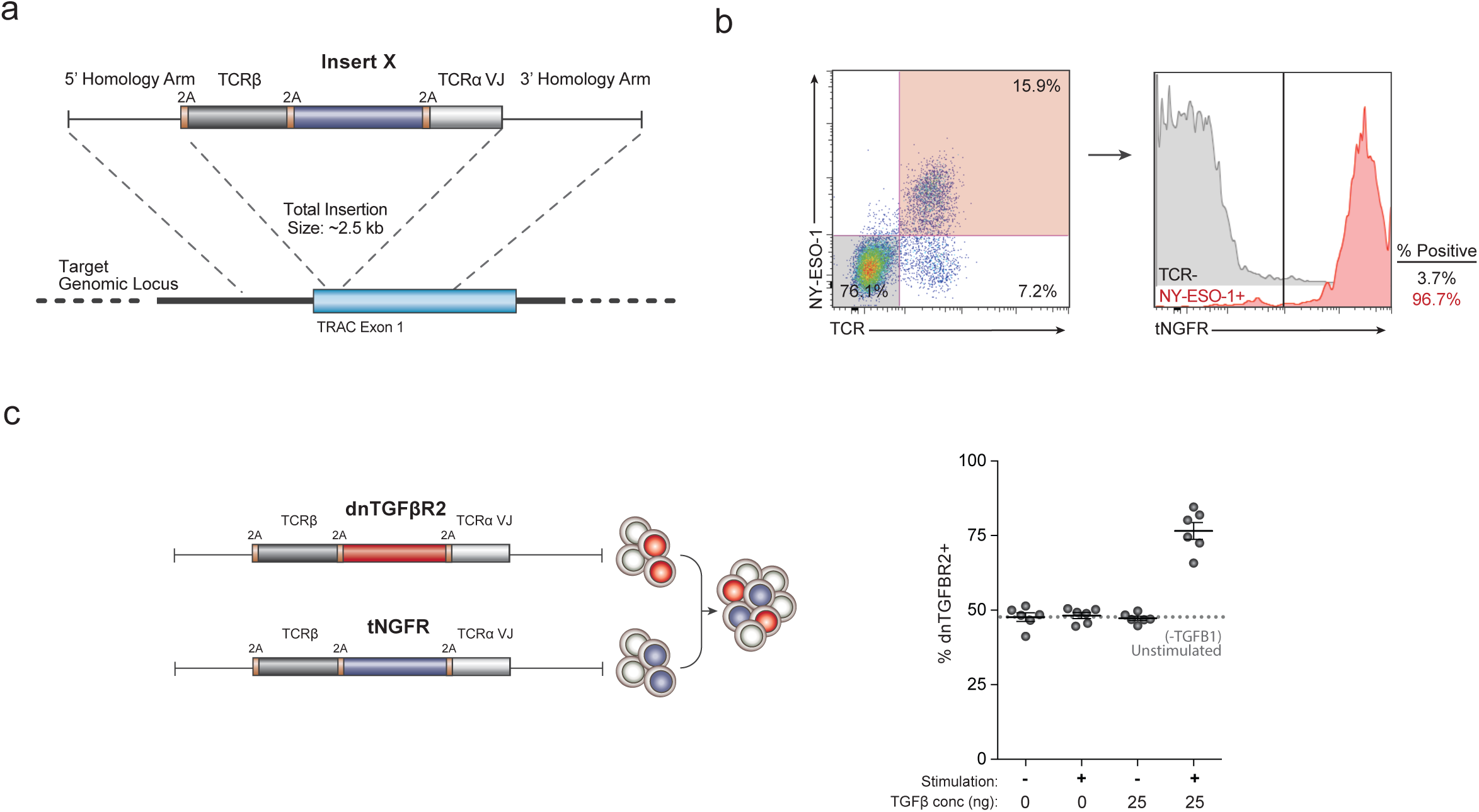
Simultaneous engineering of T cell specificity and function. **a**, Schematic of a polycistonic therapeutic knock-in cassette that enables simultaneous reprogramming of TCR specificity and integration of a co-regulated protein of interest at the endogenous TCR-α locus. The cassette includes a 2A sequence, a new full-length TCR-β chain, a second 2A sequence, the sequence for a protein of interest, a third 2A sequence, and the sequence of the new variable region of the TCR-α chain. Successful knock-in yields a polycistronic mRNA encoding new TCR-β and TCR-α chains as well as a third protein (“Insert X”). **b**, Representative data from flow cytometry assessment of a TCR+tNGFR knock-in cassette. The endogenous TCR was replaced with the NY-ESO-1 melanoma cancer antigen specific 1G4 clone and a control third protein, a truncated Nerve Growth Factor Receptor (tNGFR). Integration of this construct at the endogenous *TRAC* locus yielded NY-ESO-1 TCR+ T cells that co-expressed tNGFR. **c**, DNA sequence encoding a dominant negative TGFβ receptor 2 (dnTGFβR2)^8^ at the “Insert X” site provided resistance to the suppressive effects of TGFβ1, a cytokine found in many tumour microenvironments. NY-ESO-1 TCR+ dnTGFβR2+ T cells had a competitive expansion advantage relative to NY-ESO-1+ tNGFR+ T cells when stimulated in the presence of TGFβ1. Results are representative of n=2 healthy donors.

**Figure 3:**
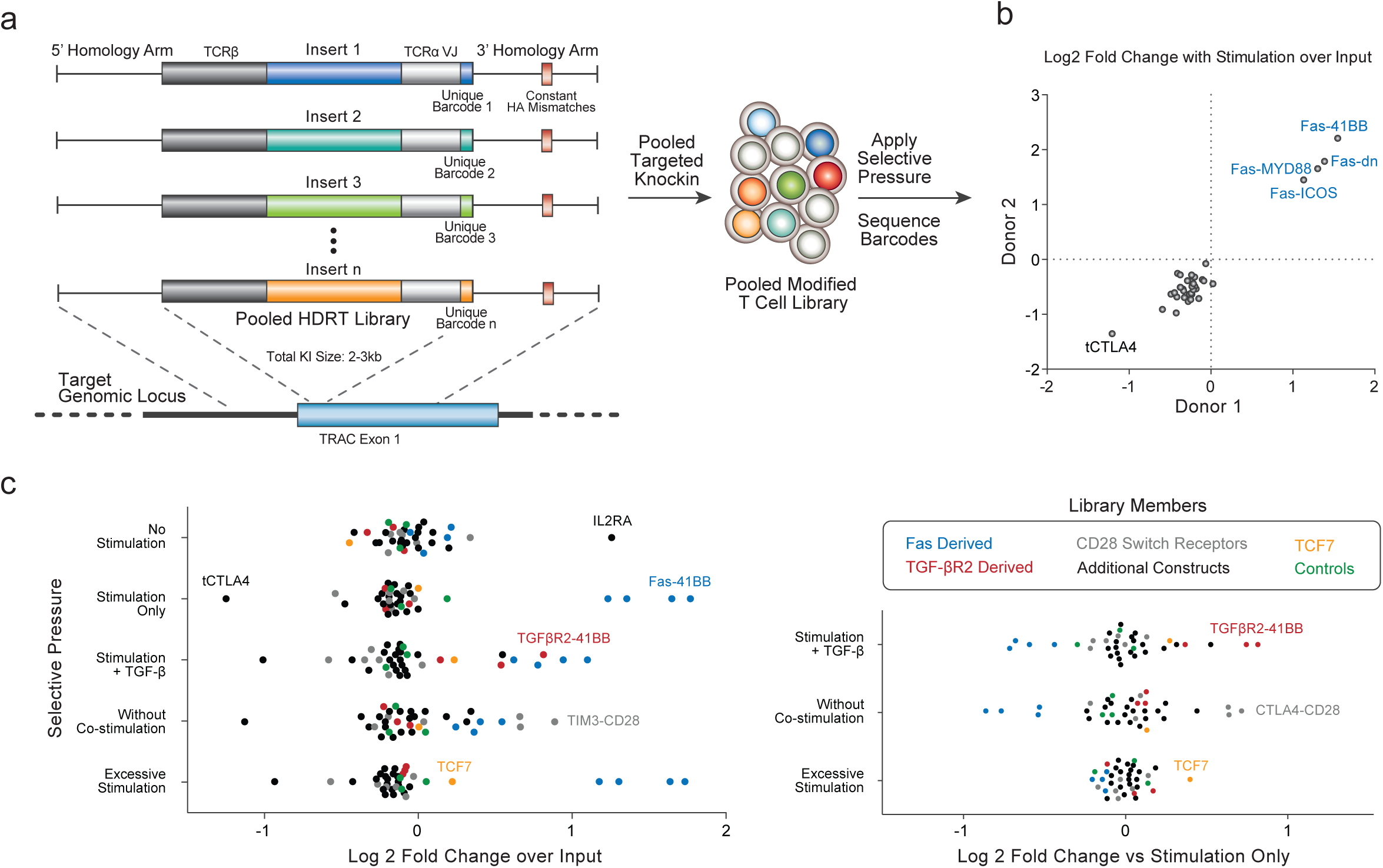
Targeted pooled knock-in screens in primary human T cells. **a**, Schematic of a generalizable method for rapid pooled knock-in screens using non-viral genome targeting. A library of HDR templates each containing a unique insert sequence and barcode is electroporated into primary human T cells to produce a heterogeneous population of modified T cells. The abundance of each knock-in modification in the cell population, after expansion with or without additional selective pressures, can be monitored by PCR and deep-sequencing of barcode sequences in each DNA insert. **b**, A 36-member pooled knock-in library was designed with dominant negative receptors, synthetic “switch” receptors with engineered intracellular domains^12^, and heterologous transcription factors, metabolic regulators and receptors (**Extended Data Fig. 13**) all targeted to the *TRAC* locus in human T cells along with a new TCR-α and TCR-β specificity for NY-ESO cancer antigen (total insert sizes ∼2-3 kB). Polycistrons encoding four different synthetic versions of apoptosis mediator FAS (blue) led to expansion of cells treated with anti-CD3/CD28 stimulation, which was reproducible across human blood donors. **c**, Assessment of pooled knock-in polycistronic *TRAC* constructs with diverse *in vitro* selective pressures of primary human T cells. Unique sets of polycistrons preferentially expanded under different selective pressures (shown relative to input library on left, and relative to CD3/CD28 stimulation on right), consistent with context-specific functions. Two novel TGFβR2 derived chimeric proteins along with the previously described dnTGFβR2 (red), drove selective expansion in the presence of exogenous TGFβ. Novel and described chimeric switch receptors of various checkpoint molecules with intracellular-CD28 (grey) drove expansion in cells treated CD3 stimulation only (no anti-CD28 co-stimulation). The polycistron encoding transcription factor TCF7 (orange) drove expansion in cells with excess TCR stimulus (5X more anti-CD3/CD28 stimulation than stimulation only condition). Averages of n=4 independent healthy donors are displayed for each condition.

**Figure 4:**
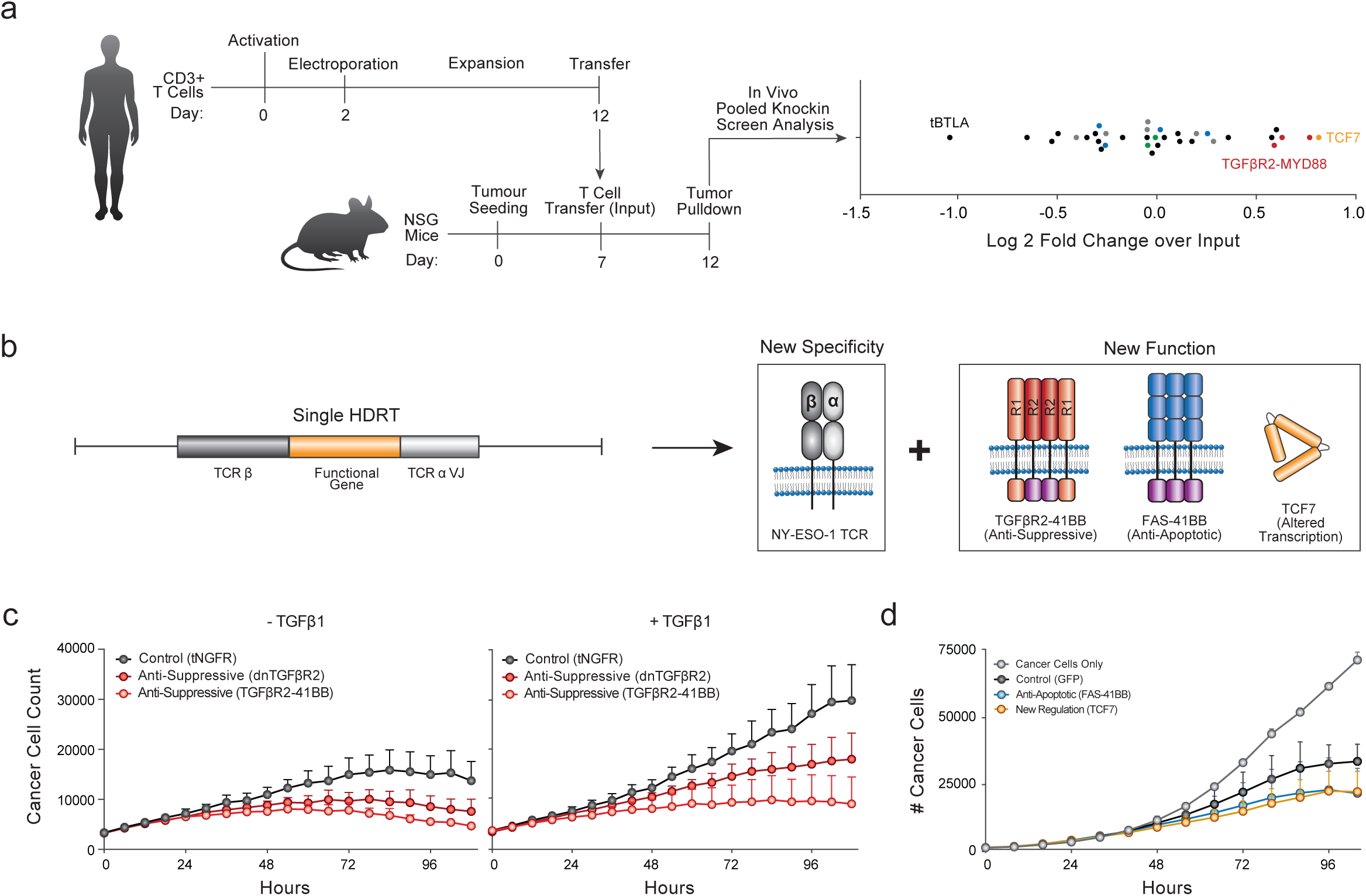
*In vivo* pooled knock-in screen and validation. **a**, *In vivo* pooled knock-in screen in a solid tumour xenograft model of human melanoma. The A375 human melanoma line expresses the target NY-ESO-1 peptide/MHC recognized by the new 1G4 TCR specificity knocked into the *TRAC* locus along with the candidate therapeutic gene constructs in polycistron library. After expansion, a bulk population of 10 million T cells (∼1 million knock-in positive NY-ESO-1 TCR+ cells) was transferred intravenously into tumour bearing mice, and an input control T cell population saved. Four days post transfer, tumours were harvested, and the modified T cell library post *in vivo* selection was sorted out and analyzed relative to input control. A subset of knock-in constructs drove T cell expansion in the *in vivo* melanoma xenograft model including polycistrons with synthetic TGFβR2 receptors and the transcription factor TCF7, which were also hits in context-specific expansion *in vitro* studies. Averages of n=2 independent healthy donors are displayed. **b**, Knock-in of a single polycistron to the *TRAC* locus allowed simultaneous replacement of the endogenous antigen specificity and co-expression of natural or synthetic gene-product to modify cell function. Complementary *in vitro* and *in vivo* pooled knock-in screening allowed rapid identification of new constructs that enhanced context-specific T cell functions, including polycistrons encoding novel TGFβR2-41BB and FAS-41BB chimeric receptors or the TCF7 transcription factor. **c**, Polycistrons with TGFβR2 switch receptor or dnTGFβR2 identified as hits in *in vitro* and *in vivo* expansion screens, enhanced NY-ESO-1+ cancer cell killing *in vitro*. A chimeric protein with TGFβR2’s extracellular domain and a 41BB intracellular domain showed greater antigen specific cancer cell killing compared to a dnTGFβR2 construct or TCR knock-in with a control tNGFR insert, both in the absence or presence of exogenous TGFβ1. **d**, Polycistrons encoding NY-ESO-1 antigen specificity plus a FAS extracellular 41BB switch receptor or the transcription factor TCF7 similarly, identified as hits in the expansion screens, showed enhanced *in vitro* NY-ESO-1+ cancer cell killing compared to TCR knock-in with a control GFP insert. Representative of n=2 independent healthy donors (**c**,**d**).

We next focused on how flexible targeted genome modifications could introduce synthetic genetic instructions at endogenous loci to modify cellular programs (**Fig. 1c**). Targeted knock-ins can integrate synthetic elements with existing sequences in a genomic locus to create Genetically Engineered Endogenous Proteins (GEEPs). We specifically wanted to test if we could engineer an endogenous gene encoding a critical receptor to reprogram: 1) gene regulation, 2) gene product, 3) extracellular domains, or 4) intracellular components (**Fig. 1c**). We show that these modifications can all be made at the *IL2RA* gene locus (**Fig. 1d-g**). We altered *IL2RA* gene regulation by knock-in of a new constitutive promoter (**Fig. 1d**) or imparted IL2RA’s endogenous stimulus-responsive expression dynamics to a new product with knock-in of a tNGFR-2A construct (**Fig. 1e**). We also separately replaced the extracellular (**Fig. 1g**) and intracellular (**Fig. 1h**) domains of IL2RA with endogenous gene modifications. GEEP T Cells could be generated across genomic loci such as *PDCD1, LAG3*, and *CD3* genes and could even be used to promote T cell expansion (**Extended Data Figs. 5-8**), demonstrating a translatable strategy to rewrite endogenous genes and gene circuits for therapeutic benefit.

With success in generating GEEP T cells, we next tested if multiple gene sequences could be integrated into a single therapeutic knock-in cassette to reprogram T cell functionality. We previously showed that the endogenous TCR-α locus could be reprogrammed with a polycistronic DNA sequence encoding a specific TCR-β and TCR-α specificity^6^. However, successful T cell therapies depend on both antigen specificity and molecular programs that govern cell fate and function. Here we tested if the polycistronic templates could be extended to include additional gene sequences that promote therapeutically-relevant cell states. First, we demonstrated that a three-gene cassette could be integrated at the endogenous TCR-α locus to both replace the endogenous TCR with a new specificity, as well as drive expression of a new gene off of the high-expression endogenous TCR promoter (**Fig. 2a**). Knock-in of a TCRβ-tNGFR-TCRα cassette to *TRAC* exon 1 showed that almost all cells with successful knock-in of the new TCR (NY-ESO-1 melanoma cancer antigen specific 1G4 clone) also showed expression of the additional tNGFR gene (**Fig. 2b**). Knock-in of a four-gene cassette to the TCR-α locus was also successful (**Extended Data Fig. 9**). *TRAC* GEEPs therefore can be engineered with co-regulated gene products.

To determine if these co-regulated gene products could enhance engineered T cell function, we replaced the tNGFR with a previously described dominant negative TGFβR2 receptor that minimizes the anti-proliferative effects of TGFβ signals^8^. Cells with NY-ESO-1 *TRAC* GEEP including dnTGFβR2 were resistant to TGFβ mediated suppression and preferentially accumulated relative to those with tNGFR when stimulated in the presence of exogenous TGFβ (**Fig. 2c** and **Extended Data Fig. 10**). Non-viral knock-in of polycistronic multi-gene program to the endogenous *TRAC* locus can successfully modify both T cell specificity and fitness.

The ability to pool multiple large sequence modifications at a specific endogenous genome locus and directly compare the functional consequences would accelerate engineering of cell therapies^9^. As a clinically relevant proof-of-concept, we hypothesized that pooled knock-in of an HDR template library, where each member contains a constant new TCR-α and TCR-β pair along with a unique third gene product, could rapidly identify new polycistronic DNA sequences that modify T cell function in therapeutically-relevant contexts (**Fig. 3a**).

Functional assessment of knock-in DNA sequences in pools depends on quantitation of the “on-target” integration of each construct in the cell population with and without selective pressures. We developed a simple deep sequencing strategy that selectively captures on-target knock-in constructs while avoiding NHEJ-edited or un-edited target genomic sequences, non-integrated HDR templates, or off-target integrations (**Extended Data Fig. 11**). Briefly, the majority of a DNA template’s homology arm sequences are not incorporated with on-target HDR integrations^10^ and a short DNA sequence that does not match the target genomic locus can be introduced into the homology arm without a large reduction in knock-in efficiency (**Extended Data Fig. 11b**,**c**). PCR primers for the inserted sequence and the genomic flanking sequence can then selectively amplify successful on-target HDR events with limited template switching (**Extended Data Figs. 11a, 12**). Addition of a barcode unique for each insert within degenerate bases of the TCR-α sequence enabled a rapid DNA sequence readout by PCR of the abundance of each individual on-target insert in the pooled population. This serves as a simple pooled knock-in methodology to target many new large DNA sequences to a specific target genomic locus and easily track their relative abundances in a cell population.

We next performed a pooled knock-in screen to discover synthetic programs that can be written into the endogenous *TRAC* locus in primary human T cells to enhance functions relevant for cancer immunotherapies. We designed a 36-member library of long DNA sequences that included published and novel dominant negative receptors, synthetic “switch” receptors with engineered intracellular domains^11,12^, and heterologous transcription factors, metabolic regulators and receptors **(Extended Data Fig. 13**). Each member of the library was cloned with self-cleaving peptide sequences between the TCRβ (full-length) and TCRα (V-region only) along with *TRAC* locus homology arms. Non-viral genome targeting with this large library of HDR templates efficiently knocked-in each library member and deep sequencing of their unique DNA barcodes accurately reflected proportions in the cell population **(Extended Data Fig. 14a-e**).

We first tested which DNA sequences in the *TRAC* locus would enhance T cell expansion upon *in vitro* stimulation. Following anti-CD3/CD28 stimulation, we observed a striking enrichment of constructs encoding various synthetic versions of the Fas receptor, especially a novel Fas switch receptor with a 4-1BB intracellular domain (**Fig. 3b** and **Extended Data Fig. 14f**). Fas is a TNF super family surface receptor that mediates apoptosis through interactions with its ligand, FASL^13^. Recently, a truncated Fas receptor was similarly shown to increase T cell persistence^14^. The results of this pooled targeted knock-in screen were highly reproducible, could be performed with earlier pooling stages, in bulk edited or sorted cells, and did not prevent robust cell expansion after electroporation **(Extended Data Fig. 14 g-k**). These data established a flexible discovery platform to identify synthetic knock-in programs that promote selectable T cell phenotypes.

Effective adoptive T cell therapies for cancer must function even in immunosuppressive environments. To identify gene modifications that enhance T cell fitness upon various challenges, we repeated the pooled screen in diverse *in vitro* conditions designed to mimic aspects of tumour micro-environments **(Fig. 3c** and **Extended Data Fig. 15a**). In the absence of restimulation, IL2RA overexpressing T cells preferentially expanded (**Extended Data Fig. 14k**), whereas with stimulation, Fas-derived synthetic receptors enabled much greater relative expansion (**Extended Data Fig. 14f**). Addition of the immunosuppressive cytokine TGFβ1 gave cells expressing the dnTGFβR2 construct a competitive advantage, and a novel chimeric TGFβR2-41BB showed even greater expansion (**Extended Data Fig. 15b**). TCR stimulation without CD28 co-stimulation selected for a variety of CD28 chimeric receptors such as a TIM3-CD28 chimera and a CTLA4-CD28 chimera (**Extended Data Fig. 15c**). Finally, excessive stimulation, which partially mirrors antigenic abundance in tumour environments, revealed a selective advantage for the transcription factor TCF7 **(Extended Data Fig. 15d**). TCF7 is a transcription factor implicated in T cells for maintenance of progenitor capacity and improved clearance in chronic viral infections^15,16^. These data from pooled knock-in screens highlight condition-specific benefits of cell states promoted by synthetic proteins or forced expression (coupled to an antigen receptor) of transcriptional regulators.

Despite the flexibility of *in vitro* assays, certain aspects of cancer immunobiology can only be assessed *in vivo*. We therefore tested if pooled knock-in screens could also be used to assess *in vivo* function of engineered human T cells. We performed an *in vivo* pooled knock-in screen using an antigen specific human melanoma xenograft model^6^. A pooled modified T cell library was transferred into immunodeficient NSG mice bearing a human melanoma expressing the NY-ESO-1 antigen, and T cells were extracted from the tumour five days later (**Fig. 4a**). Several hits from the *in vitro* screens also promoted relative *in vivo* expansion tumour environment, including TCF7 and the TGFβR2-41BB chimera (**Fig. 4a** and **Extended Data Fig. 16**). The *in vivo* screen also showed unique hits such as the metabolic protein MCT4 that had not shown enrichment in any of the *in vitro* screens performed. These results underscore the power of complementary *in vitro* and *in vivo* assays to assess synthetic gene functions. We identified three of the strongest hits and performed individual validations and *in vitro* cancer killing assays (**Fig. 4b** and **Extended Data Fig. 17**). An anti-suppressive TGFβR2-41BB chimera, an anti-apoptotic FAS-41BB chimera, and the transcriptional program altering TCF7 each improved context-dependent expansion as well as *in vitro* killing of NY-ESO-1+ cancer cells **(Fig. 4c, d** and **Extended Data Fig. 17**). Taken together, pooled knock-in screening can rapidly reveal new DNA sequences that enhance T cell functions in a variety of relevant contexts when integrated in the endogenous TCR-α locus along with a new TCR specificity.

CRISPR technology has transformed our ability to manipulate the human genome in therapeutically-relevant cell types^5–7^. High-throughput screening methods are necessary to explore the infinite number of potential manipulations possible for therapeutic relevance. We and others have made progress in genome-wide knock-out screens to discover functional pathways that can be manipulated for cell therapies^17,18^. Non-viral genome targeting allows for many more therapeutic manipulations with the addition of synthetic genetic programs. We now have developed a simple pooled knock-in screening method to discover large DNA sequences that promote cell states compatible with effective immunotherapies. Application of pooled knock-in screening *in vitro* and *in vivo* revealed novel polycistronic sequences and engineered protein products that enhanced T cell function in the challenging tumour environment in combination with reprogrammed antigen specificity. They also highlight genes that could be re-written at their endogenous loci to generate therapeutic GEEP T cells. Engineered cell therapies are an emerging “drug” class, alongside small molecules and biologics^19^. Non-viral pooled knock-in screening will accelerate the discovery and development of synthetic DNA sequences to reprogram the specificity and function of adoptive cellular therapies.

## METHODS

### Isolation of human primary T cells for gene targeting

Primary human T cells were isolated from either fresh whole blood or residuals from leukoreduction chambers after Trima Apheresis (Blood Centers of the Pacific) from healthy donors. Peripheral blood mononuclear cells (PBMCs) were isolated from whole blood samples by Ficoll centrifugation using SepMate tubes (STEMCELL, per manufacturer’s instructions). T cells were isolated from PBMCs from all cell sources by magnetic negative selection using an EasySep Human T Cell Isolation Kit (STEMCELL, per manufacturer’s instructions). Isolated T cells were either used immediately following isolation for electroporation experiments or frozen down in Bambanker freezing medium (Bulldog Bio) per manufacturer’s instructions for later use. Freshly isolated T cells were stimulated as described below. Previously frozen T cells were thawed, cultured in media without stimulation for 1 day, and then stimulated and handled as described for freshly isolated samples. Fresh blood was taken from healthy human donors under a protocol approved by the UCSF Committee on Human Research (CHR #13-11950).

### Primary human T cell culture

XVivo15 medium (STEMCELL) supplemented with 5% fetal bovine serum, 50 µM 2-mercaptoethanol, and 10 µM *N*-acetyl L-cystine was used to culture primary human T cells. In preparation for electroporation, T cells were stimulated for 2 days at a starting density of approximately 1 million cells per mL of media with anti-human CD3/CD28 Dynabeads (ThermoFisher), at a bead to cell ratio of 1:1, and cultured in XVivo15 media containing IL-2 (500 U ml^−1^; UCSF Pharmacy), IL-7 (5 ng ml^−1^; ThermoFisher), and IL-15 (5 ng ml^−1^; Life Tech). Following electroporation, T cells were cultured in XVivo15 media containing IL-2 (500 U ml^−1^) and maintained at approximately 1 million cells per mL of media. Every 2-3 days, electroporated T cells were topped up, with or without splitting, with additional media along with additional fresh IL-2 (final concentration of 500 U ml^−1^). When necessary, T cells were transferred to larger culture vessels.

### RNP production

RNPs were produced by complexing a two-component gRNA to Cas9. The two-component gRNA consisted of a crRNA and a tracrRNA, both chemically synthesized (Dharmacon or IDT) and lyophilized. Upon arrival, lyophilized RNA was resuspended in 10 mM Tris-HCL (7.4 pH) with 150 mM KCl at a concentration of 160 µM and stored in aliquots at −80□°C. Poly(L-glutamic acid) (PGA) MW 15-50 kDa (Sigma) was resuspended to 100mg/mL in water, sterile filtered, and stored in aliquots at −80C. Cas9-NLS (QB3 Macrolab) was recombinantly produced, purified, and stored at 40 µM in 20 mM HEPES-KOH, pH 7.5, 150 mM KCl, 10% glycerol, 1 mM DTT.

To produce RNPs, the crRNA and tracrRNA aliquots were thawed, mixed 1:1 by volume, and annealed by incubation at 37□ °C for 30 min to form an 80 µM gRNA solution. Next, PGA mixed with freshly-prepared gRNA at 0.8:1 volume ratio prior to complexing with Cas9 protein for final volume ratio gRNA:PGA:Cas9 of 1:0.8:1 (2:1 gRNA to Cas9 molar ratio) and incubated at 37□ °C for 15 min to form a 14.3 µM RNP solution^6,7^. RNPs were electroporated immediately after complexing.

### Double-stranded HDR DNA Template Production

Each double-stranded homology directed repair DNA template (HDRT) contained a novel/synthetic DNA insert flanked by homology arms. We used Gibson Assemblies to construct plasmids containing the HDRT sequences and then used these plasmids as templates for high-output PCR amplification (Kapa Hot Start polymerase). The resulting PCR amplicons/HDRTs were SPRI purified (1.0x) and eluted into H_2_O. The concentrations of eluted HDRTs were determined, using a 1:20 dilution, by NanoDrop and then normalized to 1 µg/µL. The size of the amplified HDRT was confirmed by gel electrophoresis in a 1.0% agarose gel. All HDR DNA template sequences used in the study are listed in **Supplementary Tables 1-3**.

### Primary T cell electroporation

For all electroporation experiments, primary T cells were prepared and cultured as described above. After stimulation for 48-56 hours, T cells were collected from their culture vessels and the anti-human CD3/anti-CD28 Dynabeads were magnetically separated from the T cells. Immediately before electroporation, de-beaded cells were centrifuged for 10 min at 90*g*, aspirated, and resuspended in the Lonza electroporation buffer P3. Each experimental condition received a range of 750,000 – 1 million activated T cells resuspended in 20 uL of P3 buffer, and all electroporation experiments were carried out in 96 well format.

For arrayed knock-in screens (**Fig. 1**), HDRTs were first aliquoted into wells of a 96-well polypropylene V-bottom plate. Poly Glutamic Acid was added between the gRNA and Cas9 complexing step during RNP assembly as described^7^. Complexed RNPs were then added to the HDRTs and allowed to incubate together at room temperature for at least 30s. For GEEPs knock-ins and pooled knock-in screens (**Fig. 2-4**), truncated Cas9 Target Sequences (tCTS) were additionally added to the 5’ and 3’ ends of the HDR template enabling a Cas9 ‘shuttle’ as described^7^. For all variations, T cells resuspended in the electroporation buffer were added to the RNP and HDRT mixture, briefly mixed, and then transferred into a 96-well electroporation cuvette plate

All electroporations were done using a Lonza 4D 96-well electroporation system with pulse code EH115. Unless otherwise indicated, 3.5 µl RNPs (comprising 50 pmol of total RNP) were electroporated, along with 1-3 µl HDR Template at 1 µg µl^−1^ (1-3 µg HDR template total). Immediately after all electroporations, 80 µl of pre-warmed media (without cytokines) was added to each well, and cells were allowed to rest for 15 min at 37□ °C in a cell culture incubator while remaining in the electroporation cuvettes. After 15 min, cells were moved to final culture vessels and media supplemented with 500 U ml^−1^ IL-2 was added.

### Arrayed Knock-in Screening

For each of 6 unique healthy donors, 5X 96 well plates of primary human T cells were electroporated. In three plates, HDR templates targeting each of 91 unique genomic loci were electroporated along with one of two on-target gRNAs or a scrambled gRNA. The final two plates were electroporated just with the on-target gRNA (complexed with Cas9 to form an RNP) but without an HDR template for amplicon sequencing. On-target and scrambled RNP plates with the HDR template were analyzed in technical duplicate for observed knock-in efficiency by flow cytometry four days following electroporation, and additionally after 24 hours of restimulation with a 1:1 CD3/CD28 dynabeads:cells ratio at five days post electroporation. Genomic DNA was isolated from the on-target gRNA only plates four days after electroporation.

After initial isolation (Day 0), immediately prior to electroporation (Day 2), and during post-electroporation expansion (Day 4), ∼1e6 CD4 and CD8 T cells from each donor were sorted by FACS for RNA-Seq and ATAC-Seq analysis (**Fig. 1b**). Half of the sorted cells were frozen in Bambanker freezing medium (Bulldog Bio) for ATAC Sequencing, and half were frozen in RNAlater (QIAGEN) for bulk RNA sequencing.

To construct an explanatory model for knock-in rates, we took a multiple linear regression approach. Briefly, this model fits the observed measured parameters with the observed knock-in rate and is described as:

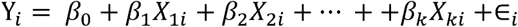

Where for the *i*th gRNA site, Y_i_ is the observed knock-in rate, β_0_ is a common intercept and Єi is the error in estimates. β_1_ to β_k_ are regression weights (or coefficients) which measure the estimate of association between the measure parameter (*X*_*ki*_) and the knock-in rate. To build the model, we averaged the measured values for across donors for each gRNA and cell type. For gene expression and chromatin accessibility, values were log transformed. The parameters used to generate the model are described in Fig 1f. The resulting model was used to explain the observed knock-in rate for all sites, across individual donors and cell types. The absolute values of all regression weights were summed and the percent of the total was determined for each parameter’s regression weight.

### Amplicon Sequencing

Genomic DNA was isolated from primary human T cells individually edited with each gRNA used in the arrayed knock-in screen in the absence of its cognate HDR template. After aspirating the supernatant, ∼100,000 cells per condition were resuspended in 20 μl of Quickextract DNA Extraction Solution (Epicenter) to a concentration of 5,000 cells per μl. Genomic DNA in Quickextract was heated to 65°C for 6 min and then 98°C for 2 min, according to the manufacturer’s protocol. 1 μl of the mixture, containing genomic DNA from 5,000 cells, was used as template in a two-step PCR amplicon sequencing approach using NEB Q5 2X Master Mix Hot Start Polymerase with the manufacturer’s recommended thermocycler conditions. After an initial 18 cycle PCR reaction with primers (**Supplementary Table 1**) amplifying an approximately 200 bp region centered on the predicted gRNA cut site, a 1.0X SPRI purification was performed and half of the samples for each biologic donor were pooled for indexing (each donor had two gRNAs that cut at each insertion site-samples for one gRNA per site were pooled, yielding two unique pools per donor). A 10-cycle PCR to append P5 and P7 Illumina sequencing adaptors and donor-specific barcodes was performed, followed again by a 1.0X SPRI purification. Concentrations were normalized across donor/gRNA indexes, samples pooled, and the library sequenced on an Illumina Mini-Seq with a 2X150 bp reads run mode.

Amplicons were processed with CRISPResso^20^, using the CRISPRessoPooled command in genome mode with default parameters. We used the hg19 human reference genome assembly. Resulting amplicon regions were matched with gRNA sites for each sample. We eliminated reads with potential sequencing errors detected as single mutated bases with no indels by CRISPResso alignment. The remaining reads were used to calculate the NHEJ percentage, or “observed cutting percentage”.

### Bulk RNA Sequencing

Total RNA from frozen samples was extracted using RNeasy Mini Kit (Qiagen) according to the manufacturer’s protocol. RNA quantification was performed using Qubit and Nanodrop 2000 and quality of the RNA was determined by the Bioanalyzer RNA 6000 Nano Kit (Agilent Technologies) for 10 random samples. We confirmed that the sample had an average RNA integrity number (RIN) that was >9 and the traces revealed characteristic size distribution of intact, non-degraded total RNA. The RNA libraries were constructed with Illumina TruSeq RNA Sample Prep Kit v2 (cat. no. RS-122-2001) according to the manufacturer’s protocol. Total RNA (500 ng) from each sample was used to establish cDNA libraries. A random set of 10 out of 36 final libraries were quality checked on the High Sensitivity DNA kit (Agilent) that revealed an average fragment size of 400bp. A total of 36 enriched libraries (3 pools of 12 uniquely indexed libraries) were constructed and sequenced using the Illumina HiSeq™ 4000 on three separate lanes at 100 bp paired end reads per sample.

RNA-Seq reads were processed with kallisto using the Homo sapiens ENSEMBL GRCh37 (hg19) cDNA reference genome annotation. Transcript counts were aggregated at the gene level. Genes of interest were subsetted from the normalized gene-level counts table and analyzed as transcripts per million (TPM).

### ATAC-Sequencing

ATAC-seq libraries were prepared following the Omni-ATAC protocol^21^. Briefly, frozen cells were thawed and stained for live cells using Ghost-Dye 710 (Tonbo Biosciences, CA, USA). 50,000 lived cells were FACS sorted and washed once with cold PBS. Technical replicates were done for most of the samples. Cell pellets were resuspended in 50µl cold ATAC-Resuspension buffer (10mM Tris-HCl (Sigmal Aldrich, MO, USA) pH 7.4, 10mM NaCl, 3mM MgCl_2_ (Sigma Aldrich,) containing 0.1% NP40 (Life Technologies, Carlsbad, CA), 0.1% Tween-20 (Sigma Aldrich) and 0.01% Digitonin (Promega, WI, USA) for 3 mins. Samples were washed once in cold resuspension buffer with 0.1% Tween 20, and centrifuged at 4C for 10 min at 300 rpm. Extracted nuclei were resuspended in 50µl of Tn5 reaction buffer (1× TD buffer (Illumina, CA, USA), 100nM Tn5 Transposase (Ilumina), 0.01% Digitonin, 0.1% Tween-20, PBS and H_2_O), and incubated at 37C for 30 min at 30 0rpm. Transposed samples were purified using MinElute PCR purification columns (Qiagen, Germany) as per manufacturer’s protocol. Purified samples were amplified and indexed using custom Nextera barcoded PCR primers as described^22^. DNA libraries were purified using MinElute columns and pooled at equal molarity. To remove primer dimers, pooled libraries were further cleaned up using AmPure beads (Beckman Coulter, CA, USA). ATAC libraries were sequenced on a Illumina HiSeq™ 4000 in paired-end 100 cycle mode.

ATAC-seq reads were trimmed using cutadapt v1.18 to remove Nextera transposase sequences, then aligned to hg19 using Bowtie2 v2.3.4.3. Low-quality reads were removed using samtools v1.9 view function (samtools view -F 1804 -f 2 -q 30 -h -b). Duplicates were removed using picard v2.18.26, then reads were converted to BED format using bedtools bamtobed function and normalized to reads per million. ATAC-seq reads mapping within a 1kb window surrounding CRISPR cut sites were counted using the bedtools intersect function.

### Flow cytometry and cell sorting

All flow cytometric analyses were performed on an Attune NxT Acoustic Focusing Cytometer (ThermoFisher). FACS was performed on the FACSAria platform (BD). Cell surface staining for flow cytometry and cell sorting was performed by pelleting and resuspending in 25 uL of FACS buffer (2% FBS in PBS) with antibodies diluted accordingly (Supplementary Table X) for 20 min at RT in the dark. Cells were washed once in FACS buffer before resuspension and analysis.

### Synthetic Product and Endogenous Product Kinetics Flow Cytometry Analysis

Non-virally edited T-cells were split into multiple replicates and analyzed by flow cytometry every day for a 5-day period starting on Day 3 after electroporation. During that 5-day period, T-cells were topped up every 2 days with additional media and IL-2, to a final concentration of 500 U/mL, with or without a 1:1 split. At Day 5 post electroporation, one set of cells was stimulated with anti-human CD3/CD28 Dynabeads and the other was left unstimulated.

### *In Vitro* Proliferation Assay

Non-virally edited T-cells were expanded in independent cultures prior to the assay. The unsorted, edited populations were pooled after approximately two weeks of expansion (with 500 U/mL of IL-2 supplemented every 2-3 days) for a competitive mixed proliferation assay.

For the CD3 competitive mixed proliferation assay, we pooled unsorted samples with CD28IC-2A-GFP, 41BBIC-2A-mCherry, or 2A-BFP knocked-in to the same CD3 complex member’s gene locus. To determine the input numbers for pooling, we accounted for the number of viable GFP+, mCherry+, or BFP+ in the respective populations (knock-in% * total viable cell count), as determined by flow cytometry analysis. The pooled sample was then distributed into round bottom 96 well plates at a starting total cell count of 50,000. The distributed samples were then cultured without stimulation, with CD3 stimulation only, with CD28 stimulation only, or with CD3/CD28 stimulation. CD3 and/or CD28 stimulation was done with plate bound antibodies. All samples were cultured in XVivo15 media supplemented with IL-2 (50 U/mL). After 4 days in culture, samples were analyzed by flow cytometry for relative outgrowth of GFP+ and mCherry+ subpopulations relative to the BFP+ subpopulation.

For the NY-ESO-1 competitive mixed proliferation assay, we pooled unsorted samples with either 1G4+dnTGFβR2+ or 1G4+tNGFR+ T cells. To determine the input number of each population, we took into account the number of viable 1G4+TCR+ in either population (knock-in% * total viable cell count), as determined by flow cytometry analysis. The pooled sample was then distributed into round bottom 96 well plates at a total starting cell count of 50,000. The distributed samples were then cultured without stimulation or with Immunocult (CD3/CD28/CD2). All samples were cultured in XVivo15 media supplemented with IL-2 (500 U/mL) with or without the addition of TGFβ1. After 5 days in culture, samples were analyzed on flow cytometry and the relative outgrowth of 1G4+dnTGFβR2+ and 1G4+tNGFR+ subpopulations was quantified.

### *In Vitro* Antigen Specific Killing Assay

A375-nRFP (NY-ESO-1^+^ HLA-A*0201^+^) melanoma cell lines stably transduced to express nuclear RFP (Zaretsky 2016 NEJM) were seeded approximately 24 h before starting the co-culture (∼1,500 cells seeded per well). Modified T cells were added at the indicated E:T ratios. The killing assay was performed in cRPMI with IL-2 and glucose. Samples were additionally topped up with TGFβ1 or an equal volume of control media. Cancer cell clearance was measured by nRFP real time imaging using an IncuCyte ZOOM (Essen) for 4-5 days and determined by the following equation: (%Confluence in A375 only wells – %Confluence in Co-culture well)/ (%Confluence in A375 only wells). At the end of the assay, cells were recovered, and the percentage of T-cells expressing various exhaustion markers was profiled by flow cytometry.

### Pooled Knock-in Screening

Targeted pooled knock-in screening was performed using the non-viral genome targeting method as described, except with ∼10bps of DNA mismatches introduced into the 3’ homology arm of the TRAC exon 1 targeting HDR template used to replace the endogenous TCR^6^. A barcode unique for each member of the knock-in library was also introduced into ∼6 degenerate bases at the 3’ end of the TCRαVJ region of the HDR template (**Fig. 4a**). The 36 constructs included in the pooled knock-in library (**Supplementary Table 3**) were designed using the Benchling DNA sequence editor, commercially synthesized as a dsDNA geneblock (IDT), and individually cloned using Gibson Assemblies into a pUC19 plasmid containing the NY-ESO-1 TCR replacement HDR sequence (except for pooled assembly conditions, whereas all geneblocks in the library were pooled prior to assembly). The library was pooled at various stages as described in figure legends (**Extended Data Fig. 12**), but unless otherwise noted HDR templates were pooled prior to electroporation.

The modified T cell libraries generated by pooled knock-ins were electroporated, cultured, and expanded as described, before being subjected to a variety of in vitro assays beginning at day 7 post electroporation and ending at day 12 post electroporation. In stimulation assays, the modified T cell library was stimulated with CD3/CD28 dynabeads at a 1:1 bead to cell ratio, and at a 5:1 bead to cell ratio for the excessive stimulation condition. In the TGFβ assay, 25 ng/mL of human TGFβ was added to the culture media. For the CD3 stimulation only condition, a 1G4 TCR (NY-ESO-1 specific) binding dextramer (Immudex) was bound to cells at 1:50 dilution in 50 uL (500,000 cells total) for 12 minutes at room temperature, prior to return to culture media. All in vitro assays began with 500,000 sorted NY-ESO-1+ T cells unless otherwise described.

At the conclusion of the in vitro or in vivo assays, T cells were pelleted and either genomic DNA was extracted (QuickExtract) or mRNA was stabilized in Trizol. mRNA was purified using a Zymo Direct-zol spin column according to manufacturer’s instructions, and converted to cDNA using a Maxima H RT First Strand cDNA Synthesis (Thermo) according to manufacturer’s instructions. Unless otherwise stated, libraries were made from isolated mRNA/cDNA. A two step PCR was performed on the isolated genomic DNA or cDNA,. The first PCR (PCR1) included a forward primer binding in the TCRαVJ region of the insert and a reverse primer binding in the genomic region overlapping the site of the mismatches in the 3’ homology arm (**Extended Data Fig. 11** and **Supplementary Table 3**), and used Kapa Hifi Hotstart polymerase for 12 cycles, followed by a 1.0X SPRI purification. The second PCR used NEB Next Ultra II Q5 polymerase for 10 cycles to append P5 and P7 Illumina sequencing adaptors and sample-specific barcodes, followed again by 1.0X SPRI purification. Normalized libraries were pooled across samples and sequenced on an Illumina Mini-Seq with a 2X150 bp reads run mode. Barcode counts from quality-filtered reads were determined in R using PDict.

### *In Vivo* Xenograft Model

An NY-ESO-1 melanoma tumour xenograft model was used as previously described^6^. All mouse experiments were completed under a UCSF Institutional Animal Care and Use Committee protocol. We used 8 to 12 week old NOD/SCID/IL-2R□-null (NSG) male mice (Jackson Laboratory) for all experiments. Mice were seeded with tumours by subcutaneous injection into a shaved right flank of 1×10^6^ A375 human melanoma cells (ATCC CRL-1619). At seven days post tumour seeding, tumour size was assessed and mice with tumour volumes between 20-40 mm^3^ were used for subsequent experiments. The length and width of the tumour was measured using electronic calipers and volume was calculated as v = 1/6 * π **\*** length * width * (length + width) / 2. Indicated numbers of T cells with the pooled knock-in library were resuspended in 100 µl of serum-free RPMI and injected retro-orbitally. A bulk edited T cell population (10×10^6^) containing at least 10% NY-ESO-1 knock-in positive cells was transferred. Five days after T cell transfer, single-cell suspensions from tumours and spleens were produced by mechanical dissociation of the tissue through a 70 µm filter, and T cells (CD45+ TCR+) were sorted from bulk tumourcytes by FACS. All animal experiments were performed in compliance with relevant ethical regulations per an approved IACUC protocol (UCSF), including a tumour size limit of 2.0 cm in any dimension.

## Data availability

Amplicon sequencing, bulk RNA sequencing, and ATAC sequencing data are available upon request. Sequencing data from pooled knock-in screens is available upon request. Plasmids containing the HDR template sequences used in the study are available through AddGene, and annotated DNA sequences for all constructs are available upon request.

## ACKNOWLEDGEMENTS

We thank members of the Marson Lab, Chris Jeans (QB3 MacroLab), Ryan Wagner and Reuben Hogan (Parnassus CAT), Vinh Nguyen (UCSF Flow Cytometry Core, P30 DK063720 and NIHS10 1S10OD021822-01) and Sarah Pyle for suggestions and assistance. T.L.R. was supported by the UCSF Medical Scientist Training Program (T32GM007618), the UCSF Endocrinology Training Grant (T32 DK007418), and the NIDDK (F30DK120213). A.M. holds a Career Award for Medical Scientists from the Burroughs Wellcome Fund, is an investigator at the Chan Zuckerberg Biohub and has received funding from the Innovative Genomics Institute (IGI), the American Endowment Foundation, a gift from Barbara Bakar, and is a member of the Parker Institute for Cancer Immunotherapy (PICI). This research was supported by NIH/NIGMS funding for the HIV Accessory & Regulatory Complexes (HARC) Center (P50 GM082250; AM)

## AUTHOR INFORMATION

### Contributions

T.L.R. designed and performed arrayed knock-in experiments. T.L.R., P.J.L., R.A., A.T., R.Y., and J.H. designed and cloned arrayed knock-in constructs. T.L.R. and R.A. performed amplicon sequencing. T.L.R. and M.L.Y.N. performed ATAC sequencing. T.L.R. and Y.L. performed RNA sequencing. T.L.R., R.Y., D.W. and E.S. performed computational analysis of arrayed knock-in data. T.L.R. and P.J.L, designed GEEPs constructs and performed GEEPs experiments. T.L.R. designed the pooled knock-in strategy. T.L.R., P.J.L. and J.N. designed constructs for pooled knock-ins. T.L.R. and J.N. performed pooled knock-in experiments. T.L.R., J.N. and E.S. computationally analyzed pooled knock-in experiments. D.N.N. advised on the use of anionic polymers. D.G., J.A.B. and K.R. advised on the development of pooled knock-in screening. T.L.R., P.J.L. and A.M. wrote the manuscript with input from all authors.

### Competing Financial Interests

The authors declare competing financial interests: T.L.R. and A.M. are co-founders of Arsenal Therapeutics. A.M. is a co-founder of Spotlight Therapeutics. A.M. and J.A.B. are co-founders of Sonoma Biotherapeutics. A.M. serves as on the scientific advisory board of PACT Pharma and was a former advisor to Juno Therapeutics. J.A.B. is a consultant for Juno, a Celgene company; a stock holder and member of the Board of Directors of Provention and Rheos Medicines; a member of the Scientific Advisory Board of Pfizer Center for Therapeutic Innovation; a member of the Scientific Advisory Board and stock holder for Vir Therapeutics, Arcus Biotherapeutics, Quentis Therapeutics, Solid Biosciences, NeoCept (Founder) and Celsius Therapeutics (Founder). JAB owns stock in MacroGenics Inc., Provention, Viacyte Inc., and Kadmon Holdings. The Marson Laboratory has received sponsored research support from Juno Therapeutics, Epinomics, Sanofi and a gift from Gilead. Patents have been filed based on the findings described here.

### Corresponding Author

Correspondence and requests for materials should be addressed to.

## EXTENDED DATA FIGURES AND TABLES

**Extended Data Figure 1: Large non-viral knock-ins at 91 genomic loci in primary human T cells.**

**a**, A non-viral arrayed knock-in screen was performed across 91 unique genomic loci. Efficient knock-in of a GFP fusion protein to the C terminus of TCR-α and the four members of the CD3 complex were all achieved. No HDR template control showed minimal background levels of fluorescence in the GFP channel.

**b**, For additional targets, a tNGFR-2A polycistronic cassette or GFP fusion was knocked in to the N-terminus or C-terminus of each target gene. Efficient knock-in was achieved at most of 24 additional surface receptor target genes. In both GFP fusion constructs and tNGFR-2A targeted constructs the observed GFP or tNGFR expression was driven by each gene’s endogenous promoter, yielding diverse expression levels across target loci. For example, note the high expression of tNGFR targeted to the *B2M* or *CD45 (PTPRC)* loci, and the lower expression at *CXCR4*. No knock-in was observed at some target sites, such as *CX3CR1* and *LTK*, whereas over 50% of cells were successfully targeted at other sites, such as *IL2RA* and *CD28*.

**c**, Cells with various checkpoint receptor genes targeted showed greater knock-in reporter expression following re-stimulation (**Extended Data Fig. 2**). Note, all cells received an initial anti-CD3/CD28 activation upon isolation and two days prior to electroporation in order to achieve efficient non-viral genome targeting, and flow cytometry was performed either four days after electroporation without additional stimulation (“Resting”) or five days after electroporation following 24 hours of anti-CD3/CD28 bead stimulation (**Extended Data Fig. 2**).

**d**, Non-viral genome targeting at 16 additional transcription factor genes. Some target loci, such as *JunD*, showed low observed knock-in percentages but high expression levels of the knock-in reporter, whereas other sites, such as *NCOA3*, showed high percentages of observed knock-in but low knock-in reporter expression levels.

**e**, Efficient targeting of seven additional genes encoding cytoskeleton components. Again note the variable expression levels of the knock-in reporters under control of different endogenous promoters.

**f**, Non-viral knock-in results for an additional 32 target genes.

All displays are from the same healthy blood donor, and are representative of n=6 total donors tested during the arrayed knock-in screen. Displays show the more efficient of the two gRNAs tested for each locus. Unless significant differences in knock-in % were observed between CD8 and CD4 T cells or between re-stimulated and resting conditions (**Extended Data Fig. 2**), the resting CD8 T cell condition is shown. Panels showing re-stimulated cells are indicated with red plus signs; panels showing CD4 data are labeled in parentheses. In all panels the X-axis is either GFP fluorescence or tNGFR staining, and the Y-axis shows cell size (FSC-A).

**Extended Data Figure 2: Analysis of observed knock-in percentages across 91 target loci in multiple cell types and stimulation conditions.**

**a**, Arrayed knock-in screen of large HDR templates (either GFP or tNGFR, ∼800 bps) across 91 unique genomic target sites, testing two gRNAs per sites in both CD4+ and CD8+ T cells in 6 unique healthy human donors.

**b**, Arrayed knock-in screen readouts. Observed knock-in percentages were analyzed at the cellular level by flow cytometry for GFP or tNGFR reporter expression (day 6 or 7). To determine contribution of target gene expression to observed knock-in, RNA-Seq was performed at day 0 (prior to activation), day 2 (time of electroporation), and day 4 (during expansion). To determine contribution of target site accessibility, ATAC-Seq was performed at days 0 and 2. Amplicon sequencing to determine the actual cutting efficiency of each RNP was performed at day 6 (using separate RNP only plates where no HDR template was electroporated).

**c**, Relative observed knock-in percentages in CD8 vs CD4 T cells. The highest divergence in observed knock-in between the cell types was for genes encoding their hallmark surface receptors, *CD8A* and *CD4* respectively. Percentge of cell expressing knock-in reporter at *41BB* (TNFRSF9) and *LAG3* was higher in CD8 T cells, while observed percentage at *IL2* was higher in CD4 T cells. The majority of targeted sites did not show large differences between the two cell types. Observed knock-in % for n=6 donors across 91 target genomic loci with 2 gRNAs per locus.

**d**, Relative observed knock-in percentages in re-stimulated vs resting CD8 T cells. The amount of knock-in observed by flow cytometry at various activation/exhaustion markers, such as *PDCD1, 41BB (TNFRSF9)*, and *OX40* (*TNFRSF4*) was higher after a second stimulation four days following electroporation. In comparison, observed knock-in at other sites, such as *FBL, CCR2*, and *IL7R (CD127)*, was higher without a second stimulation (“Unstimulated”). n=6 donors across 91 target genomic loci with 2 gRNAs per locus.

**e**, Analysis of the observed off-target knock-in % for each of the 91 unique HDR templates containing a GFP or tNGFR knock-in reporter sequence along with homology arms specific for their target genomic locus. In all 6 donors in the arrayed knock-in screen, all 91 HDR templates were electroporated with a scrambled gRNA (forming an RNP designed not to target any specific site in the human genome). While the majority of HDR templates showed minimal observed off-target knock-in, a handful of HDR templates (targeting the genes *FBL, IL2RG*, and *STAT2*) showed higher amounts. Future analysis of the DNA sequences of these templates could yield further insights into patterns of off-target integration.

**f**, Observed MFI of knock-in positive cells across all templates, donors, and cell types, was correlated (*R*^2^ = 0.50) with the RNA-Seq expression values recorded for each combination of target gene, donor, and cell type. Aggregated data from n=6 unique human blood donors.

**g**, Predicted on-target cut score^23^ for each gRNA used in the arrayed knock-in screen (91 target sites × 2 gRNA per site = 182 total gRNAs) was weakly correlated (*R*^2^ = 0.03) with the observed editing efficiency in each of the 6 donors tested. All 182 gRNAs were individually electroporated into bulk CD3+ T cells in all 6 donors in the absence of an HDR template, and the % editing at each target locus was analyzed by amplicon sequencing. Likely due to the high efficiency of RNP based knock outs in primary human T cells (vast majority of gRNAs showed >95% NHEJ editing by amplicon sequencing), the predicted cut score was not strongly correlated with observed editing in these conditions.

**Extended Data Figure 3: Correlation of gRNA and target DNA locus parameters with observed knock-in efficiency.**

**a**, The distance between the cut site of the tested gRNAs and the integration site of their associated HDR template in bps (“Cut Distance”) was weakly negatively correlated with observed knock-in efficiency across all donors. The necessity of short distances between a cut site and integration site has been well described^24^, but within the window of a cut distance less than approximately 25 bps there was a low correlation with observed knock-in.

**b**, A gRNA can recognize a DNA sequence and cut in either the 5’ or 3’ direction relative to the integration site. A cut towards the 5’ direction was defined as when the gRNA’s NGG PAM faced towards the integration site in a 5’ to 3’ direction, and was assigned a value of −1. A cut towards the 3’ direction was defined as when the gRNA’s NGG PAM faced away from the integration site in a 5’ to 3’ direction, and was assigned a value of 1. Directionality of the cut was not clearly correlated with observed knock-in rates across the 91 targeted loci.

**c**, The predicted on-target cut score^23^ for each guide was not clearly correlated with observed on-target knock-in percentage.

**d**, The observed editing rate (NHEJ) of each gRNA in each of the 6 donors tested (**Extended Data Fig. 2e**) showed a weak positive correlation (*R*^2^ = 0.059) with observed knock-in efficiency. X-axis displays proportion of alleles with NHEJ edit.

**e**, Bulk RNA-Seq was performed in all 6 healthy donors tested for 2 cell types (CD4 and CD8 T cells) at three time points. Expression levels of the 91 target genes at the time of T cell isolation and prior to activation (“day 0”), at the time of electroporation two days after anti-CD3/CD28 stimulation (“day 2”), or during the expansion phase after electroporation (“day 4”) were determined. RNA expression levels at all three time points were correlated with observed knock-in %, with the highest correlation (*R*^2^ = 0.28) being the time point (day 4) closest to the time of the protein level flow cytometry readout. Note that the expression level of the reporter for each construct in the arrayed knock-in screen is driven by the target gene’s endogenous promoter, so observed knock-in rates assessed by flow cytometry are confounded by expression of the target gene. Genes that are expressed at levels below the detection limit of the flow cytometric readout could potentially have higher actual knock-in percentages that are not observed due to a low level of protein expression. X-axis displays log10 transcripts per million (TPM).

**f**, ATAC-Seq was performed in all 6 tested healthy donors for 2 cell types (CD4 and CD8 T cells) at two time points (Day 0 before activation and Day 2 prior to electroporation). DNA accessibility was determined for a 1 kb window centered on the cut site of each gRNA at the 91 target loci. At both timepoints, the accessibility of the target locus was correlated (*R*^2^ = 0.18 on Day 2) with observed knock-in efficiency. X-axis displays log10 reads per million (RPM).

**g**, A multivariate linear regression model (**Fig 1e, f**) incorporating each of the gRNA parameters (except predicted cutting), RNA expression, and DNA accessibility showed greater correlation (*R*^2^ = 0.39) than any individual parameter in isolation.

**Extended Data Figure 4: Examination of knock-in target sites with divergent predicted and observed knock-in efficiencies.**

**a**, Analysis of difference between the predicted knock-in efficiencies by a multivariate linear regression model and the observed knock-in efficiencies for each of the 91 unique genomic target sites with 2 gRNAs per site in 2 cell types (CD4 and CD8 T cells). The majority of genes had predicted knock-in efficiencies within a one-fold change of the observed amount, but a handful of genes had much higher predicted knock-in efficiencies than were observed (*ELOB, JUND*) and some genes had much lower predicted knock-in values than were observed (*DDX20, STAT4, ITGB1*).

**b**, Analysis of 6 gene targets with higher predicted knock-in % than observed. The two tested gRNAs are colored, and the two lines for each guide represent CD4 and CD8 T cells.

**c**, Analysis of 6 gene targets with lower predicted knock-in % than observed. As these sites showed higher knock-in efficiencies than would otherwise be predicted, further examination of these targets and their sequence context may reveal design features that could improve overall knock-in efficiencies across target sites.

**d**, Analysis of 6 target loci with the highest variance in prediction accuracy between the two gRNAs tested at that site. For at least two of these sites (*SATB1, CCR7*) the gRNAs with higher predicted knock-in than was observed were found to cut the associated HDR template due to design errors in the DNA HDRT sequence (the gRNA binding sequence and/or PAM site for all gRNAs should be altered in their respective HDR template to prevent cutting of the HDR template either prior to integration or in the genome after integration).

**e**, Analysis of the top 6 target loci with the highest variance in prediction accuracy between the two cell types tested (CD4 and CD8 T cells).

Averages from n=6 unique healthy donors are displayed (**a-e**).

**Extended Data Figure 5: Genetically Engineered Endogenous Proteins (GEEPs) with synthetic regulation of endogenous products.**

**a**, Schematic representation of knock-in strategy for targeting a novel promoter to a gene of interest with or without a selectable reporter (tNGFR).

**b**, To test whether we could reprogram endogenous gene regulation, we targeted an SFFV promoter to the 5’ non-coding region of *PDCD1* (PD1) or LAG3 in addition to *IL2RA* (CD25) (shown here are results from a second human blood donor), all well-established stimulation-responsive immune receptors. When cultured without re-stimulation for 7-11 days following electroporation, T cells with the SFFV promoter knock-in (On-Target RNP) showed sustained, stimulation-independent expression of PD1, LAG3, and IL2RA when analyzed by flow cytometry. In contrast, T cells without SFFV promoter knock-in (Control RNP) showed baseline expression levels for each corresponding protein.

**c**, Representative flow cytometry data of edited T cells where we integrated (in 5’ to 3’ order) an SFFV promoter, a selectable reporter (tNGFR), and a 2A sequence such that a polycistronic mRNA driven by the SFFV promoter encodes both tNGFR and the endogenous protein. We targeted the N-terminus of three immune receptors, PD1, LAG3, and IL2RA, whose expression is induced upon T-cell stimulation (Top). We observed low expression levels of each immune receptor in cells that have been cultured for 7 days post-electroporation without re-stimulation. Consistently, we observe that in control conditions (Scrambled RNP + HDR DNA Template) expression levels of immune receptor are relatively low. In the on-target knock-in conditions (On-target RNP + HDR DNA Template), we see that tNGFR+ cells, which also have the SFFV promoter targeted with the same template, have high expression levels of the respective immune receptor, while the tNGFR-cells have expression levels similar or lower than the control, the latter most likely attributed to knockout occurring with the on-target RNP in the absence of HDR. When we re-stimulated these cells, expression levels of the immune receptors increased in the control samples. In the re-stimulated on-target samples, the tNGFR+ cells retained high expression levels of the respective immune receptors, whereas the tNGFR-cells showed induced expression levels, although to a lesser extent than controls (likely due to gene knockout).

**d**, Comparison of tNGFR reporter expression levels with expression levels of the targeted immune receptors in resting control and on-target knock-in treated cells confirmed that on-target cells have high expression levels of both tNGFR and the targeted endogenous immune receptor (demonstrated by the linear relationship), while the control cells have lower expression levels of the respective immune receptors and negligible tNGFR expression.

**e**, Having validated our knock-in strategy for integrating a novel/synthetic promoter along with a selection marker, we applied the knock-in strategy to an array of transcription factors whose overexpression may be beneficial for T cell proliferation and long-term function. We knocked in tNGFR to transcription factor loci with 2A sequences to preserve expression of their endogenous gene products. To readout successful integration of our construct, we assessed tNGFR expression in on-target samples for four different transcription factors and found that we were able to achieve 10-25% knock-in efficiency. This strategy has implications for being able to efficiently modulate transcription factor expression to influence T-cell fate and function.

All displays are representative of n=2 independent healthy donors.

**Extended Data Figure 6: Creation of Genetically Engineered Endogenous Proteins (GEEPs) at the *PDCD1* locus.**

**a**, Schematic representation of knock-in strategy for targeting novel protein(s) to the N-terminus of a gene of interest for either: (1) coordinated expression of the novel protein(s) and the endogenous protein (inclusion of 2A sequence), or (2) expression of the novel protein(s) under endogenous gene regulation with knock out of the endogenous protein (inclusion of a polyA termination signal).

**b**, Representative flow cytometry data validating our strategy for coordinated expression of a novel protein and PD1 under the endogenous gene regulation of *PDCD1*. In resting cells (top row), there is minimal PD1 and tNGFR expression. However, by 48 hours after re-stimulation with anti-CD3/CD28 Dynabeads, we see a coordinated induction of both tNGFR and PD1.

**c**, Representative flow cytometry data validating our strategy for expression of a novel protein under the endogenous gene regulation of *PDCD1* with knockout of endogenous PD1. In resting cells (top row), there is minimal PD1 and tNGFR expression. However, by 48 hours after re-stimulation with anti-CD3/CD28 Dynabeads, we see induction of tNGFR without PD1.

**d**, Schematic representation of knock-in strategy for altering the extracellular receptor domain at an endogenous gene locus. To replace the extracellular domain of PD1, we targeted a 2A-NGFR Extracellular Domain construct to Exon 3 of *PDCD1*, which encodes the PD1 transmembrane domain. The 2A sequence should ensure that endogenous extracellular receptor is cleaved from protein product and not expressed on the cell surface.

**e**, Representative flow cytometry data validating our strategy for replacing extracellular receptor domain. In resting T-cells (top row), there is minimal PD1 expression and NGFR staining. Upon restimulation with anti-CD3/CD28 Dynabeads, we observed an NGFR+PD1-T cell population. The chimeric NGFR-PD1 receptor is programmed into the *PDCD1* gene locus with endogenous stimulation-responsive gene regulation.

All displays are representative of n=2 independent healthy donors.

**Extended Data Figure 7: Genetically Engineered Endogenous Proteins (GEEPs) with endogenous regulation of synthetic products.**

**a**, Schematic representation of knock-in strategy for targeting a novel protein to the N-terminus of a gene of interest with a 2A excision element for coordinated expression of the novel protein and the endogenous protein under endogenous gene regulation.

**b**, Representative flow cytometry data from experiments where we integrated a tNGFR-2A construct at the N-terminus of *IL2RA*. We demonstrate tNGFR expression levels mirrored stimulation-dependent regulation of the endogenous targeted protein. In cells where *IL2RA* was targeted, we observed a linear relationship between IL2RA and tNGFR (with high expression of both) at day 3 post-electroporation, indicative of coordinated expression of the two proteins. At day 7 post-electroporation, cells that were cultured without re-stimulation had a gradual and coordinated decrease in expression of both IL2RA and tNGFR, whereas a IL2RA high and tNGFR high population was induced in re-stimulated cells. MFI of tNGFR (left) and IL2RA (right) are quantified across 7 days post-electroporation in resting and re-stimulated CD4 and CD8 T cells.

**c**, Representative flow cytometry data from experiments where we integrated a tNGFR-2A construct at the N-terminus of CD28. We observed a linear relationship between CD28 and tNGFR (with high expression of both) at day 3 post-electroporation, indicative of coordinated expression of the two proteins. CD28 and tNGFR expression levels both remained high without re-stimulation at day 7. Upon re-stimulated, we observed a simultaneous negative modulation of CD28 and tNGFR expression. The more pronounced decrease of CD28 expression could be due to the combination of gene expression modulation and internalization of the protein, whereas tNGFR should not be internalized upon stimulation.

**d**, Representative flow cytometry data from experiments where we integrated a tNGFR-2A construct at the N-terminus of *LAG3*. At Day 3, LAG3 and tNGFR expression levels were low and both levels of expression remained low without restimulation at Day 7. Upon re-stimulation, we observed the simultaneous induction of LAG3 and tNGFR by day 7.

All displays are representative of n=2 independent healthy donors.

**Extended Data Figure 8: Creation of Genetically Engineered Endogenous Proteins (GEEPs) with synthetic signaling in CD3 complex members.**

**a**, Schematic representation of three different constructs we designed to modify the C-terminus of each CD3 subunits in the TCR complex, which include the CD3δ chain, CD3ε chain, CD3γ chain, and CD3ζ chain. For initial tests, we designed a construct that would knock-in a 2A-BFP at the C-terminus of each of the different CD3 subunit gene. The 2A-BFP integration would create a polycistronic mRNA that produces two separate proteins: an unmodified CD3 chain and BFP. Once the 2A-BFP integration was validated, we modified the construct to include a cytoplasmic domain of an co-stimulatory immune receptor (CD28 or 41BB) before the 2A sequence (followed by GFP or mCherry) such that the C-terminus of the CD3 subunit chain now contains an additional signaling domain/motif.

**b**, To readout successful integration of the signaling domain, we analyzed the percentage of fluorescent protein expressing T cells by flow cytometry. The addition of a co-stimulatory signaling domain did not have a significant/consistent effect on knock-in efficiency. Although the positioning of the additional signaling domain relative to endogenous CD3 signaling motifs was not optimized, the ability to modify the intracellular domains of individual CD3 subunits provides a promising platform for tuning TCR signaling.

**c**, Results of competitive mixed culture assays testing the growth advantage conferred by synthetic intracellar signaling domains on CD3 components. We pooled unsorted edited T cells with CD28IC-2A-GFP, 41BBIC-2A-mCherry, or 2A-BFP knocked-in to the same CD3 complex member’s gene locus. We then cultured the mixed cell population without stimulation, with CD3 stimulation only, with CD28 stimulation only, or with CD3/CD28 stimulation. After 4 days in culture, samples were analyzed by flow cytometry for relative outgrowth of GFP+ or mCherry+ subpopulations relative to the BFP+ subpopulation. We then normalized the proportions to those found in the corresponding unstimulated condition.

All displays are representative of n=2 independent healthy donors.

**Extended Data Figure 9: Knock-in of a four-component polycistronic cassette to the human *TRAC* locus.**

**a**, Schematic representation of the strategy for simultaneous in-frame integration of a new replacement TCR specificity and two additional co-regulated proteins of interest at the endogenous TCR-α locus. We designed a single HDR DNA Template that included (in order) a Furin-Spacer-T2A sequence, the sequence for a new full-length TCR-β chain, a Furin-Spacer-E2A sequence, the sequence for our first protein of interest, a Furin-Spacer-F2A sequence, the sequence for our second protein of interest, a Furin-Spacer-P2A sequence, and the sequence of the new variable region of the TCR-α chain. These exogenous sequences were flanked by arms homologous to the endogenous *TRAC* locus Exon 1 region. Successful knock-in would yield a polycistronic mRNA that encodes four separate proteins.

**b**, Representative flow cytometry data of T cells with knock in of a model four-component polycistronic cassette. For initial tests, the endogenous polyclonal TCR repertoires was re-written with the 1G4 TCR (recognizes NY-ESO-1 cancer antigen) and our additional proteins of interest were tNGFR and GFP. Proper integration of this construct at the endogenous *TRAC* locus would yield NY-ESO-1 TCR+ tNGFR+ GFP+ T cells. The flow cytometry plot in the top left illustrates the knock-in efficiency, determined by the percentage T cells staining positive with a NY-ESO-1 dextramer. NY-ESO-1+ cells almost all express GFP and tNGFR concordantly (top right flow plot) whereas NY-ESO-1-TCR-cells largely do not (bottom left flow plot). A relatively small percentage of TCR+ NY-ESO-1-cells express both GFP and tNGFR, but not either alone (bottom right flow plot). This observation can most likely be explained by off-target integration of our construct at a locus with active expression or an on-target integration of our construct with improper TCR expression or an on-target integration of the cassette with TCR mis-pairing of either the 1G4 TCR-α chain, TCR-β chain, or both with endogenous TCR components.

All displays are representative of n=2 independent healthy donors.

**Extended Data Figure 10: Characterization of T cell function after knock-in of a new TCR specificity along with co-regulated dnTGF**β**R2.**

**a**, Our TCR+Payload knock-in strategy at the endogenous *TRAC* locus leads to coordinated gene expression of a novel TCR-β chain, a defined TCR-α chain, and a protein of interest. The three proteins, however, should retain independent post-translational regulation. To test this, we sorted NYESO-1+ tNGFR+ T cells and compared NY-ESO-1 TCR expression levels with tNGFR expression levels in resting cells versus after TCR re-stimulation. TCR stimulation is known to cause TCR internalization, and after 24 hours of stimulation, we observed decreased NY-ESO-1 TCR expression by dextramer staining. In contrast, tNGFR protein expression remained high after 24 hours of stimulation as expected.

**b**, Histogram shows gating strategy to determine relative expansion of NY-ESO-1 TCR+ dnTGFβR2+ T-cells over NY-ESO-1 TCR+ tNGFR+ in an *in vitro* competitive growth assay. The majority of T cells at this stage of the experiment (19 days after initial isolation, 2 rounds of stimulation, continuous culture in 500 U/mL of IL-2) were CD8+ T cells. Thus, we focused our flow analysis only on CD8+ T cells. Gating on NY-ESO-1+CD3+ CD8+ T cells, we observed a bimodal distribution of cells when examining tNGFR expression. The percentage of tNGFR-NY-ESO-1+CD3+ CD8+ T cells was inferred to represent the NY-ESO-1 TCR+ dnTGFβR2+ T cells for downstream analyses. **c**, The results of a replicate *in vitro* competitive growth assay in another independent human blood donor. After 5 days, we again found an expansion of the NY-ESO-1 TCR+ dnTGFβR2+ T cells relative to the NY-ESO-1 TCR+ tNGFR+ T cells in stimulated pooled samples cultured in 25 ng/mL of TGFβ1.

All displays are representative of n=2 independent healthy donors.

**Extended Data Figure 11: Simple DNA sequencing strategy to selectively detect on-target knock-ins.**

**a**, DNA sequencing of HDR outcomes is complicated by the large amount of HDRT introduced into cells that remains unintegrated. A successful on-target knock-in must be distinguished from the wild type or NHEJ modified genomic locus, non-integrated template, and homology-independent off-target integrations. To overcome these challenges, two aspects of HDR were used to create a unique amplifiable sequence exclusively at on-target knock-ins. First, only a short region of the homology arms of an HDRT are integrated into the genome during HDR (along with the entire length of the inserted region), while the majority of the homology arms are used for complementary base pairing with the targeted genomic locus. Second, small mismatches in the homology arms can be tolerated during HDR, as long as the vast majority of the homology arms remain complementary to the genomic target site. This enabled a strategy where a short stretch of mismatches was introduced to a homology arm (∼10 bp of mismatches to the 3’ HA in this case). This mismatched sequences will be found in all non-integrated templated and in homology-independent off-target integrations (as the entire homology arms should be integrated during NHEJ-mediated integrations at off-target sites of random dsDNA breaks). However, at the on-target HDR-mediated integrations, the mismatches in the homology arm should not be integrated into the genome. This enables a simple PCR to amplify the on-target locus using one primer complementary to the inserted sequence (and thus unable to prime off genomic locus without knock-in), and a second primer complementary to the genomic sequence at the sequence where the HDR homology arm has a mismatch. Only the on-target genomic knock-in sites should possess both primer binding sites and these will be selectively amplified.

**b**, Knock-in of a polycistronic sequence to the *TRAC* locus replacing the endogenous TCR with a new specificity (NY-ESO-1 antigen) along with an additional co-regulated reporter gene (tNGFR) with standard homology arms (left) or with a 3’ HA containing ∼10 bp of mismatches to the target genomic site ∼100 bps from the insert sequence (right). Succesful HDR (assessed by tNGFR reporter expression and NY-ESO-1 TCR dextramer staining on flow cytometry) was still achieved even with the homology arm mismatch template.

**c**, Quantification showing that knock-in of ∼2.5kb NY-ESO-1 TCR+tNGFR was slightly less efficient with the homology arm mismatches compared to unaltered homology arms, but still easily detectable. Two technical replicates in n=3 independent

**Extended Data Figure 12: Analysis of template switching with varying pooling stages in pooled knock-in screens.**

**a**, Pooling of samples can occur at each distinct step of a non-viral genome targeting protocol: dsDNA fragments containing the unique members of a pooled knock-in library can be pooled prior to assembly into DNA plasmids already containing constant elements such as homology arms (“Pooled Assembly”); DNA plasmids containing the entire HDRT sequence for each unique library member can be pooled prior to a PCR reaction to generate large amounts of dsDNA HDR template (“Pooled PCR”); dsDNA HDR templates for each unique library member can be pooled prior to electroporation into the final cells (“Pooled Electroporation”); or, cells separately electroporated with each unique library member can be pooled following electroporation but before a final readout (“Pooled Culture”).

**b**, A library of two members, GFP and RFP encoded within a polycistronic knock-in cassette encoding a new TCR specificity (NY-ESO-1 specific 1G4 clone) to *TRAC* exon 1, was used to analyze template switching rates with different stages of pooling. Knock-in positive primary human T cell could be identified based on expression of the new TCR specificity (TCR+ NY-ESO-1+ assessed by dextramer staining). Successful knock-in was observed with all pooling strategies.

**c**, Knock-in positive cells were analyzed for GFP and RFP expression by flow cytometry. Cells with either GFP or RFP templates alone only showed expression of each respective fluorescent protein, while the Pooled Culture condition showed equal populations of GFP and RFP positive cells exclusively, without GFP+RFP+ cells. Pooling conditions prior to the electroporation step (Pooled Assembly, Pooled PCR, or Pooled Electroporation) all showed both single GFP+ or RFP+ cells, as well as dual GFP+RFP+ cells, potentially due to bi-allelic knock-in at the *TRAC* locus, as T cells can express TCR-α chains off of both alleles^6^. Multiple populations were sorted for barcode sequencing, including bulk knock-in negative cells (NY-ESO-1-), bulk knock-in positive cells (NY-ESO-1+), and individual populations of RFP+GFP- or RFP-GFP+ cells. Next-generation DNA sequencing of on-target knock-ins was performed using either isolated mRNA converted to cDNA, or isolated genomic DNA using a 2 step PCR. An initial PCR amplified the HDR-integrated barcode region using a reverse primer overlapping mismatches in the 3’ HA of the HDR template (**Extended Data Fig. 11**) and a constant forward primer within the insert sequence (total amplified region ∼140bp). A second indexing PCR was then performed prior to pooling of samples for sequencing.

**d**, To analyze the selectivity of the on-target knock-in PCR sequencing strategy, the total amount of amplification from sorted knock-in positive (NY-ESO-1+) vs knock-in negative (NY-ESO-1-) cells was analyzed relative to the bulk population of edited cells using a constant amount of input genomic DNA prior to the first PCR and quantifying the total relative number of reads sequenced (no concentration normalizations were used between samples at any protocol steps). Knock-in positive cells showed preferential amplification of the knocked-in sequence with the barcode relative to the bulk edited population, while knock-in negative cells showed almost no successful amplification, consistent with expected selectivity for amplifying and sequencing on-target knock-ins relative to non-integrated HDRT sequence or off-target integrations (**Extended Data Fig. 11**).

**e**, The degree to which alleles without a knock-in were amplified during the barcode sequencing PCR was analyzed across pooling stages and comparing isolated cDNA vs genomic DNA (gDNA). All conditions showed low amounts of reads without a barcode sequence (e.g. containing the wild-type sequence at the genomic locus), although when sequencing off of cDNA the amount was consistently slightly higher (∼1% of total reads). Sequencing off of cDNA has the advantage of amplifying the number of barcodes from each individual cell, but requires the pooled knock-in screen be performed in a coding region that is expressed (such as the *TRAC* locus) and that the barcode be integrated into degenerate bases in a coding sequence. In contrast, sequencing amplified genomic DNA has the advantage of generalizability to any genomic locus where a successful knock-in can be performed (**Figure 1**), but has potentially lower signal to noise compared to sequencing off of mRNA (converted to cDNA) when using low numbers of cells.

**f**, The percentage of sequenced reads (from either genomic DNA or cDNA amplification) that contained the GFP HDR template’s barcode corresponded with the observed percentage of cells expressing GFP protein by flow cytometry across pooling conditions. This confirmed the ability of the pooled knock-in screening sequencing strategy to accurately assess the knock-in frequencies in a mixed population by sequencing of DNA barcodes.

**g**, The percentage of sequenced barcodes in sorted GFP+ or RFP+ cells that contained the correct barcode is displayed across pooling conditions when sequencing from amplified genomic DNA. Knock-in of GFP or RFP templates only yielded 100% of reads containing the correct barcode, and pooled culture of cells after electroporation yielded >99% correct barcodes. However, pooling at earlier experimental stages produced a consistent increased amount of template switching across donors in either GFP+ (left) or RFP+ (right) sorted cells.

**h**, The frequency of template switching using the homology arm mismatch priming strategy for pooled knock-in screening was quantified across pooling stages. The amount of template switching observed was consistent between sequencing from amplified genomic DNA or cDNA. The earliest pooling stage, Pooled Assembly, showed the greatest amount of template switching, but a consistent amount of template switching was observed with Pooled PCR and Pooled Electroporation conditions, indicating that crossing over or template switching events likely occurred during the Gibson Assembly reaction, the PCR to produce the HDR templates, and potentially even within the cell. Given that in a pooled knock-in library with two members (GFP and RFP) approximately half of the actual amount of template switching will yield a barcode with an identical sequence, the predicted amount of template switching in an arbitrarily large library will be higher (**Supplementary Note 1**). Given the parameters of the current pooled knock-in library design (∼400 bps between unique library insert and its corresponding barcode, separated by the new knocked in TCR-α specificity), the amount of predicted template switching with pooled assembly reactions was ∼50%, whereas with a pooled electroporation was only ∼10%. Given the relatively low rate of template switching with the pooled electroporation with the constructs tested here, we decided to move forward with this pooling strategy for functional screens.

All experiments display one representative donor (**b, c**) or one or more technical replicates (**d-h**) from n=2 unique healthy donors.

**Extended Data Figure 13: Design of a 36-member pooled knock-in library to alter T cell function.**

**a**, A pooled knock-in library of 36 potentially therapeutic genes was constructed that could be integrated along with a new TCR specificity (NY-ESO-1) using a single HDR template (**Supplemetary Table 3**). The library was designed to contain both previously published and novel members that potentially modified immuno-therapeutic T cell function in a variety of broad classes: immune checkpoints with their intracellular domains either truncated (“tPD1” or “tCTLA4”) or replaced with an “activating” domain (chimeric switch receptors, “CTLA4-CD28”); apoptotic mediators similarly truncated or with intracellular domains switched; genes involved in cell proliferation; chemokines; transcription factors; genes involved in metabolic pathways associated with survival in tumour environments; and suppressive cytokine receptors either as truncated/dominant negative receptors (“dnTGFβR2”) or with intracellular domains switched.

**b**, All 36 constructs were synthesized and placed into a TCR insertion cassette that would replace the endogenous T cell receptor with a new specificity (NY-ESO-1 TCR) as well as drive expression of the new gene that potentially modifies T cell function with endogenous TCR-α gene regulation. Each library member was individually tested in an arrayed knock-in screen and assayed for the percent knock-in as well MFI of the surface expressed TCR to assay any potential effects of the individual inserts on TCR expression in cells from two human blood donors.

**c**, All 36 constructs successfully showed functional TCR expression as analyzed by surface dextramer staining for the new NY-ESO-1 TCR.

**d**, The total insert sizes ranged from ∼2,000-3,000 bps (not including the homology arm sequences), and little correlation was observed between template size and knock-in efficiency.

**e**, Consistent expression levels of knock-in TCR (assessed as MFI of NY-ESO-1 TCR) were observed following targeting of all 36 library members individually.

Complete design strategy, amino acid sequences, and DNA sequences for all 36 members of the library are contained in **Supplementary Table 3**.

**Extended Data Fig. 14: Technical validations of pooled knock-in screening in primary human T cells.**

**a**, Pooled knock-in screening of a 36-member HDR template library where each member encodes the same antigen specificity (NY-ESO-1 specific TCR) as well as a unique gene with barcode that potentially modifies T cell function, all targeted for integration at the *TRAC* locus. After electroporation, a modified T cell population is generated that can then be assayed, for instance by addition of a second TCR stimulation (an initial stimulation is required to knock-in the constructs). The frequency of the each barcode present in the population can be determined by DNA sequencing. Barcode frequencies can then be compared to the input population to see the relative effects of each library member on T cell behavior in that assay.

**b**, Two constructs in the 36-member library were easily detectable by flow cytometry: controls encoding GFP and RFP. Gating on knock-in positive cells that have acquired the new NY-ESO-1 specific TCR revealed that the proportion of cells that were also GFP+ or RFP+ was roughly equivalent.

**c**, Distribution of sequence barcodes in the modified T cell library seven days after pooled electroporation of the 36-member library. The percentage of total reads for each library member was consistent across four unique healthy human T cell donors, and the library showed a relatively even distribution (Gini coefficient = 0.048).

**d**, Good correspondence was found between observed population frequencies at the protein level by flow cytometry and detected barcode frequencies at the DNA level through the pooled knock-in sequencing approach. For GFP and RFP constructs, which were easily observable by flow cytometry, the proportion of cells positive at the protein level was similar to the proportion of reads with corresponding GFP and RFP barcodes.

**e**, Plot shows the relationship between the size of the inserted sequence and the detected frequency in the population of modified T cells. The full-length NY-ESO-1 TCRβ and NY-ESO-1 TCRα VJ segments along with their associated 2A elements are ∼1.5 kb, while the size of the additional functional gene knocked in in the same construct varied from ∼0.5 – 1.5 kb, yielding a total insert size of ∼2 – 3 kb. A weak negative correlation was observed with larger inserts present in the library at slightly lower frequencies (*R*^2^ = 0.11).

**f**, Seven days after pooled electroporation of the 36 pooled knock-in constructs, the modified T cell population was either stimulated anti-CD3/CD28 (1:1 beads:cells ratio) or isolated as an input population. The log2 fold change in barcode frequency after 5 days of in-vitro TCR stimulation is displayed relative to the input population. Constructs derived from the apoptotic mediator FAS cell surface protein showed notable increases in relative frequency across four unique healthy T cell donors demonstrating that they confer a fitness advantage to T cells under these conditions.

**g**, The reproducibility of pooled knock-in screen results was examined across technical replicates and for different pooling stages (**Extended Data Fig. 12a**). Technical replicates of the TCR stimulation screen in the same biologic donor showed strong correlation (*R*^2^ = 0.99). The correlation between Pooled Assembly and Pooled Electroporation conditions was also strong, albeit less so (*R*^2^ = 0.88). This could be due to greater variation between technical replicates in the Pooled Assembly condition from more frequent template switching observed when the library pooling occurs at earlier stages (**Extended Data Fig. 12h**), as the correlation between technical replicates of Pooled Assembly conditions was similarly slightly weaker (*R*^2^ = 0.89).

**h**, The number of knock-in positive, viable cells is important for pooled screens. The expansion of primary human T cells after pooled knock-in was assayed for 10 days post electroporation. Given 1 million primary human T cells at isolation, an average of ∼0.5 million knock-in positive cells were recovered by four days post-electroporation (average overall knock-in efficiencies were ∼10% – 20%), and these cells continued to expand robustly over additional days in culture without re-stimulation across four healthy human donors.

**i**, Knock-in experiments generate mixed populations of cells, some with alleles containing the desired knock-in, some with knockout alleles, and some with unedited alleles (**Extended Data Fig. 14b**). Pooled knock-in screening can be performed on both sorted knock-in positive cells (here sorted on NY-ESO-1 dextramer stained cells) as well as an unsorted bulk population of targeted cells when sorting is not practical or feasible. The sequenced barcode frequencies after pooled knock-ins were highly correlated between sorted and unsorted bulk populations (*R*^2^ = 0.87).

**j-k**, For the majority of pooled knock-in experiments, T cells were expanded for 7-10 days without re-stimulation after electroporation prior to application of a selective pressure. Expansion in culture (containing media + IL-2 only) over this time period did not show any large changes in abundance of library members, except for a large relative increase in abundance of the IL2RA-encoding construct (averages of four donors shown in **Fig. 3c**).

Experiments display or are representative of n=2 (**d, g, i**) or n=4 (**c, e-f, h, j-k**) unique healthy human T cell donors. Dotted lines (**k**) represent maximum and minimum abundance of control library members (encoding GFP, RFP and tNGFR).

**Extended Data Fig. 15: Pooled knock-in screening identifies distinct functional sequences under varying *in vitro* selective pressures mimicking aspects of tumour environments.**

**a**, Pooled electroporation of a 36-member library of DNA sequences encoding potential function modifying proteins along with a defined TCR specificity (NY-ESO-1 antigen) generated a pooled library of modified primary human T cells. Various *in vitro* selective pressures mimicking aspects tumour environments could then be applied and the distribution of unique barcodes in the population of modified T cells could be compared: 1) to the input population of T cells, or 2) among the selective pressures. These analyses revealed which constructs conferred competitive fitness advantages to T cells in each specific context.

**b**, Distribution of library construct abundances after *in vitro* culture for 5 days in TGFβ, represented as a ranked list of log2 fold changes over the input library population. Input cells were taken at 7 days post electroporation and anti-CD3/CD28 (1:1 beads:cells) stimulation was applied along with 25 ng/mL of exogenous TGFβ in the culture media. Relative to input, multiple FAS derived anti-apoptotic receptors as well as TGFBR2 derived anti-suppressive receptors increased relative expansion. When compared to anti-CD3/CD28 bead-based stimulation alone, Fas derived receptors showed a relative decrease in abundance (but still an absolute increase) demonstrating potentially enhanced susceptibility to TGFβ-mediated suppression. TGFβR2 derived receptors in contrast showed by far the greatest relative expansion in the presence of TGFβ. The previously published dominant negative TGFBR2 receptor (dnTGFBR2)^8^ and a novel chimeric TGFβR2 (41BB intracellular domain) switch receptor both drove strong T cell expansion in TGFβ.

**c**, Stimulation of the T cell population modified with library through the TCR only (through incubation with an NY-ESO-1 specific dextramer) without the presence of a CD28-engaging co-stimulatory signal showed selective increased abundance of some, but not all, CD28 chimeric switch receptors. The extracellular domains of various immune checkpoint proteins, such as CTLA4, TIM3, and BTLA were fused with an intracellular domain of CD28. Constructs encoding CTLA4-CD28, TIM3-CD28, and BTLA-CD28 conferred selective advantage to cells cultures with TCR stimulation alone, relative to those cells cultured with anti-CD3/CD28 stimulation.

**d**, In the context of excessive amounts of TCR stimulation (anti-CD3/CD28 5:1 bead:cell ratio instead of a standard 1:1 ratio), Fas receptor-derived constructs again showed increased relative abundance when compared to the input population. When comparing the suppressive excessive stimulation population to standard stimulation, the Fas constructs again showed greater relative inhibition in the excessive stimulation, whereas a construct expressing the transcription factor TCF7 conferred a greater relative advantage in cells with excessive anti-CD3/CD28 stimulation (when compared to standard anti-CD3/CD28 stimulation) in all four blood donors.All graphs display log 2 fold change compared to modified T cell library input, or relative log 2 fold change compared to CD3/CD28 stimulation. Mean of n=4 unique healthy donors is displayed and was used to rank the constructs. Dotted lines represent max and min abundance of non-functional control library members.

**Extended Data Fig. 16: *In vivo* pooled knock-in screen in humanized mouse solid tumour xenograft model.**

**a**, After generation and expansion for 10 days *in vitro*, a modified population of human T cells electroporated with the 36-member *TRAC* polycistronic knock-in library (2.5e6 sorted NY-ESO-1 dextramer stain positive T cells) was adoptively transferred into immunodeficient NSG mice bearing a human melanoma tumour xenograft (A375 melanoma cells expressing the target peptide/MHC for the NY-ESO-1 TCR) injected subcutaneously 7 days before transfer. After 5 days of *in vivo* selective pressure in the solid tumour environment the tumours were dissected, T cells were sorted, and the abundance of barcodes relative to input population was analyzed by DNA sequencing.

**b**, Biologic replicates of the *in vivo* solid tumour pooled knock-in screen with cells from different human blood donors showed greater variance across the library than *in vitro* pooled knock-in screens (**Fig. 4b**), but consistently showed the same top library hits.

**c**, Technical replicates of the *in vivo* pooled knock-in screen with cells from the same blood donor similarly showed greater variance than *in vitro* pooled knock-in screens (**Extended Data Fig. 14g**).

**d**, Multiple hits from *in vitro* pooled knock-in screens also drove T cell expansion *in vivo* in the solid tumour xenograft environment. The construct encoding the transcription factor TCF7 and constructs encoding TGFβR2 derived chimeric receptors showed robust and reproducible increases in relative abundance *in vivo*. Additional library members not identified as hits in any of the *in vitro* screens performed, such as the metabolite transporter MCT4, showed strong relative enrichment *in vivo* in the tumour environment.

Experiments display or are representative of n=2 (**b-d**) unique healthy human T cell donors. Dotted lines represent maximum and minimum abundance of control library members (encoding GFP, RFP and tNGFR).

**Extended Data Fig. 17: Individual validation of hits from pooled knock-in screening with *in vitro* NY-ESO-1+ cancer cell killing assay.**

**a**, Individual functional validation of a TGFβR2-41BB chimeric receptor bearing the extracellular domain of the suppressive cytokine receptor TGFβR2 and the intracellular domain of the proliferative receptor 41BB. With a polycistronic HDR template, primary human T cells were engineered to express both a new TCR specificity (NY-ESO-1) as well as the TGFβR2-41BB chimeric switch receptor.

**b**, In the presence of TGFβ, the TGFβR2-41BB modified cells recapitulated the observed phenotype of greater relative expansion compared to stimulation only (**Extended Data Fig. 15**). Sorted NY-ESO-1+ T cells also expressing either TGFβR2-41BB or a GFP control were re-stimulated with anti-CD3/CD28 beads (1:1 bead to cell ratio) 7 days after electroporation and expansion was assayed by quantifying absolute cell counts at each indicated day. Surface staining for activation and exhaustion markers was performed 6 days after the stimulation.

**c**, TGFβR2-41BB modified cells showed greater NY-ESO-1+ cancer cell killing *in vitro* than tNGFR controls, and comparable if not greater killing than the dnTGFβR2 modified cells, when co-cultured with A375 human melanoma cells with the addition of exogenous TGFβ across the indicated range of T cell to cancer cell ratios. At 5 days after beginning the co-culture killing assay, T cells were removed and stained for surface expression of PD1.

**d**, Individual functional validation of a Fas-41BB chimeric receptor bearing the extracellular domain of the apoptotic receptor FAS and an intracellular domain of the proliferative receptor 41BB. With a polycistronic HDR template, primary human T cells were engineered to express both a new TCR specificity (NY-ESO-1 antigen) as well as the chimeric Fas-41BB receptor.

**e**, Expression of a Fas-41BB chimeric receptor increased relative expansion compared to expression of a GFP control receptor (both along with the new TCR specificity) in an antigen-independent proliferation assay (anti-CD3/CD28 bead re-stimulation 7 days post electroporation), validating the observed increased expansion seen with stimulation in the pooled screens (**Fig. 4c**). Crucially, increased expansion with the Fas-41BB receptor was only seen upon re-stimulation, whereas continued expansion in IL-2 without re-stimulation showed no relative expansion advantage compared to control. Decreased surface expression of some activation and exhaustion markers was also observed after bead stimulation.

**f**, T cells targeted with the NY-ESO-1 TCR / Fas-41BB construct showed greater NY-ESO-1+ cancer cell killing *in vitro* than those targeted with control NY-ESO-1 TCR construct.

**g**, Individual functional validation of the TCF7 expression construct. With a polycistronic HDR template, primary human T cells were engineered to express both a new TCR specificity (NY-ESO-1 antigen) as well as an altered transcriptional program through TCF7 controlled by endogenous TCR-α gene regulation.

**h**, Expression of TCF7 recapitulated the higher observed relative expansion compared to NY-ESO-1 TCR+ GFP+ control knock-in under excessive stimulation conditions (5:1 anti-CD3/CD28 bead to cell ratio) relative to standard stimulation (1:1 bead to cell ratio). Expression of the indicated activation and exhaustion markers did not appear changed between the modifications.

**i**, T cells targeted with the NY-ESO-1 TCR / TCF7 construct showed greater NY-ESO-1+ cancer cell killing *in vitro* than those targeted with control NY-ESO-1 TCR construct.

Experiments display or are representative of n=2 (**b-c, e-f, h-i**) unique healthy human T cell donors.

## SUPPLEMENTARY INFORMATION

**Supplementary Table 1**

Sequence and design information from arrayed knock-in screen (**Fig. 1**) for all 91 target genomic sites, including gRNA sequences and DNA HDRT sequences.

**Supplementary Table 2**

Sequence and design information from all GEEPs constructs (**Fig. 2, 3**), including gRNA sequences and DNA HDRT sequences.

**Supplementary Table 3**

Sequence and design information from 36-member library of polycistronic constructs used in pooled knock-in screens (**Fig. 4**), including gRNA sequences and DNA HDRT sequences.

**Supplementary Table 4**

A list of antibodies used in this study.

**Supplementary Note 1**

Calculation of predicted total amount of template switching with an arbitrarily large library size given observed amounts of template switching from a two-member library.

**Supplementary Note 1: Calculation of predicted total template switching for a large library using observed template switching from a two-member library.**

An estimation of the total amount of template switching (barcode switching) during a large pooled experiment where there is some distance between a DNA barcode and the functional DNA sequence with which the barcode is associated can be inferred using the observed amount of template switching from a small library (**Extended Data Fig. 12**). A simple model requires two assumptions:

*Assumption 1*: Template switching is random, and any barcode that is swapped is equally likely to switch to any other barcode.

*Assumption 2*: A template that has switched once is equally likely to switch again.

Using a two-member library where DNA barcodes are associated with observable functional DNA sequences (e.g. encoding a detectable surface marker or fluorescent protein), cells that are confirmed to express the functional DNA sequence can be isolated from all other cells that express the other (e.g. sorting for GFP+ cells) and their DNA barcodes sequenced. The percentage of the sequenced DNA barcodes that does not match the barcode associated with the functional DNA sequence is thus the observed amount of template switching, represented as a ratio.

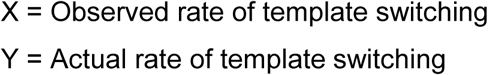

In a two-member library half of the actual template switching will occur with another template that contains the same barcode, and is thus not detected. Therefore, the observed amount of switching must be multiplied by 2.

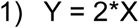

However a template that has switched once is equally likely to switch again, and therefore additional rounds of switching after the first can occur.

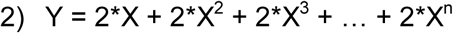

Simplifying the equation yields:

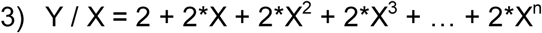

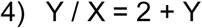

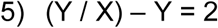

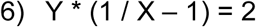

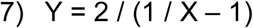

Thus, a general equation for total template switching (Y) can be derived from the observed amount of template switching (X) in a two-member library. For example, if the observed template switching rate was 0.2 (20% of total reads contained an incorrect barcode), then the actual predicted rate of template switching would be 0.5 (50% of the reads are predicted to have switched, but because many switched to a template containing the same barcode, they were not observed). In a library of arbitrarily large size, there will be no significant switching with templates containing the same barcode, and thus the observed and actual amounts of template switching will converge.

