## Extended Data Figures 1-17 for "Rapid discovery of synthetic DNA sequences to rewrite endogenous T cell circuits"

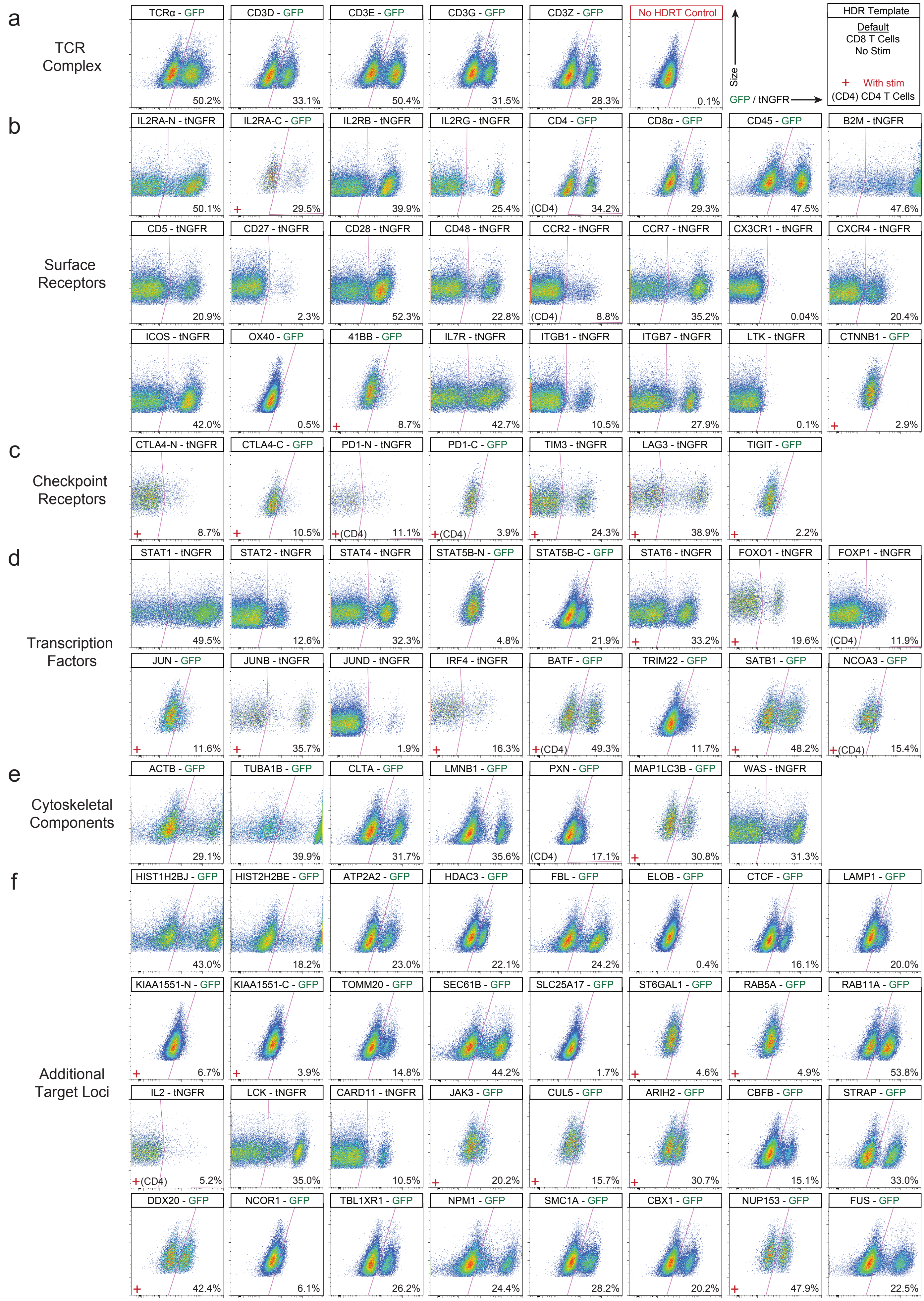


Extended Data Fig. 1

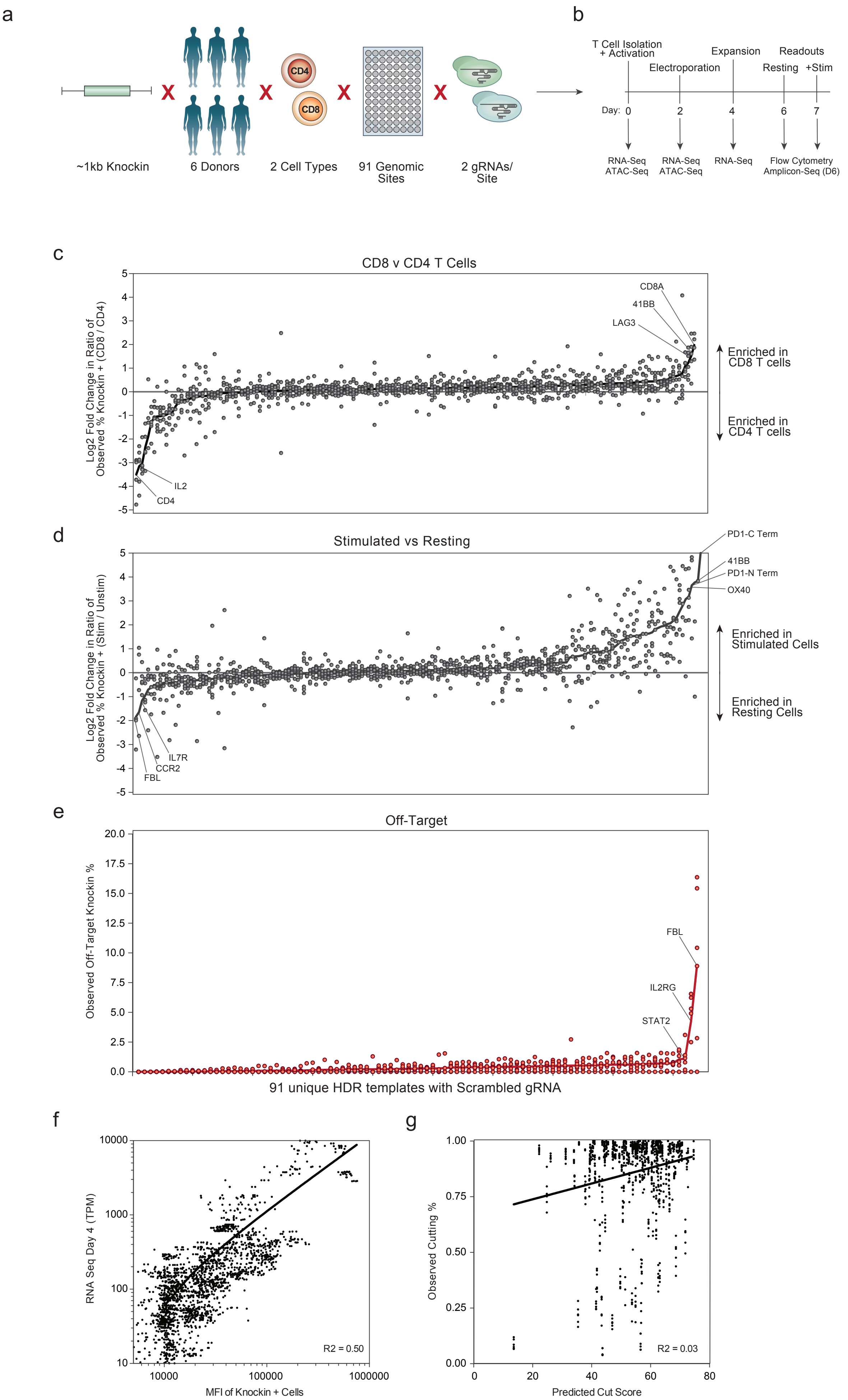


Extended Data Fig. 2

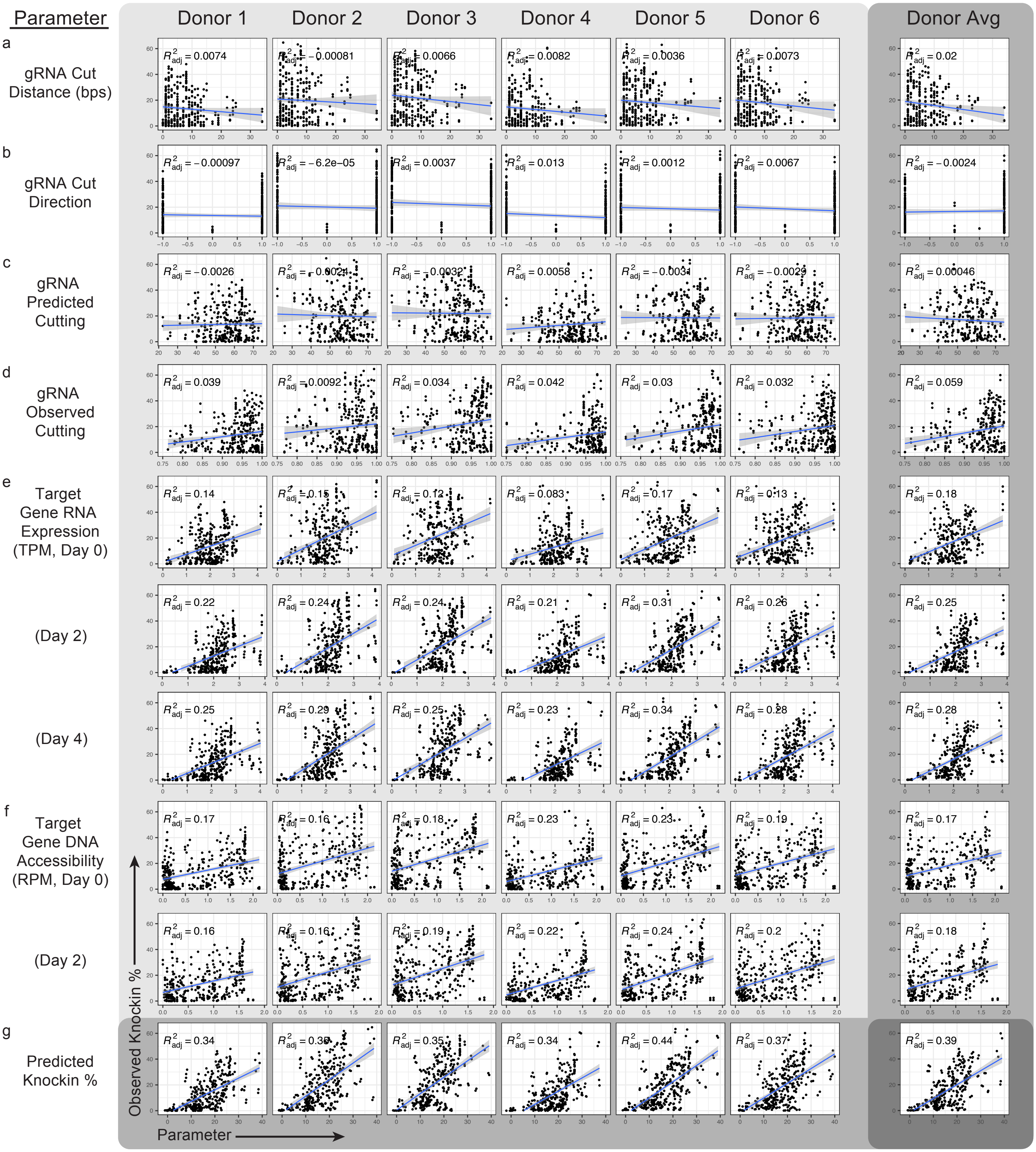


Extended Data Fig. 3

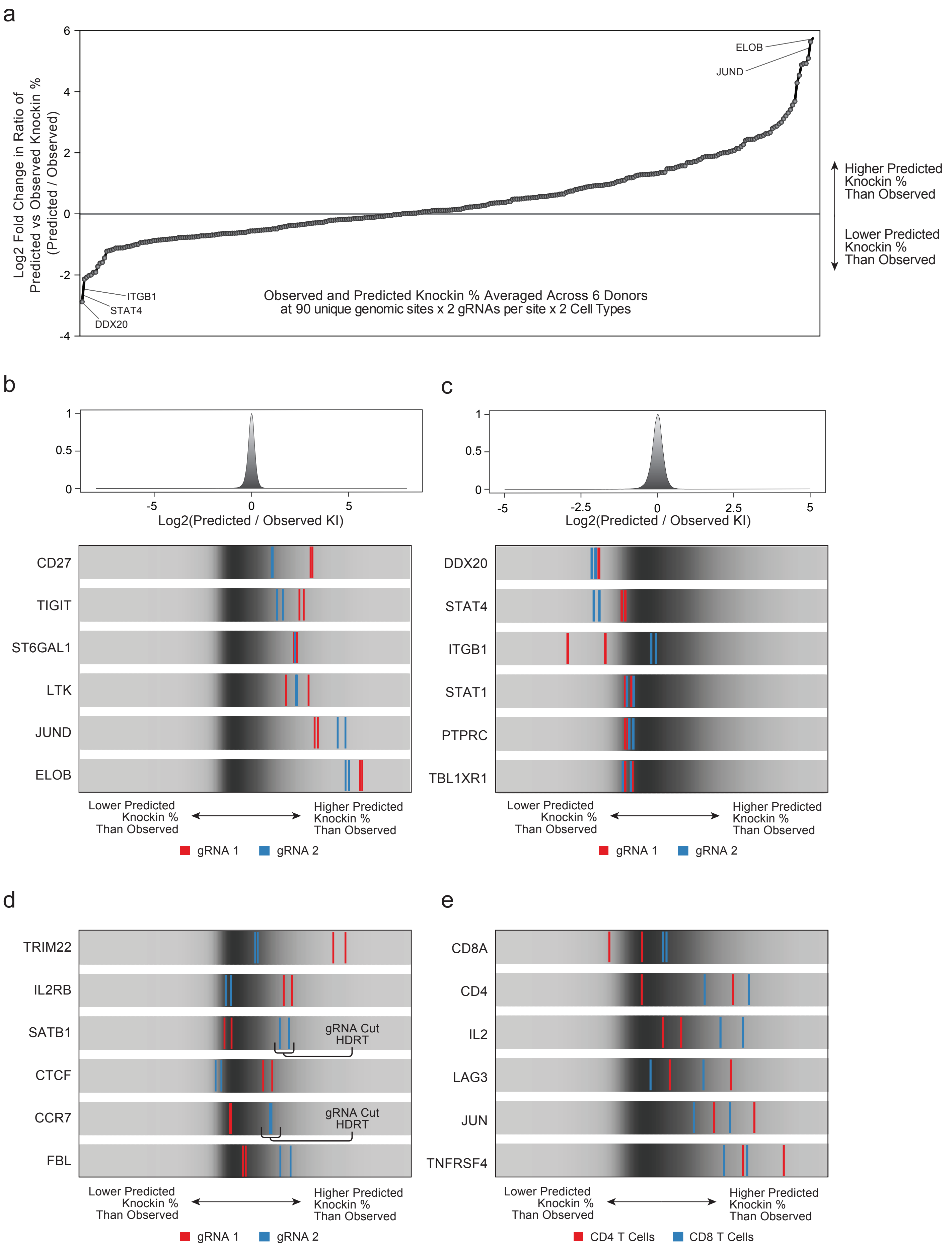


Extended Data Fig. 4

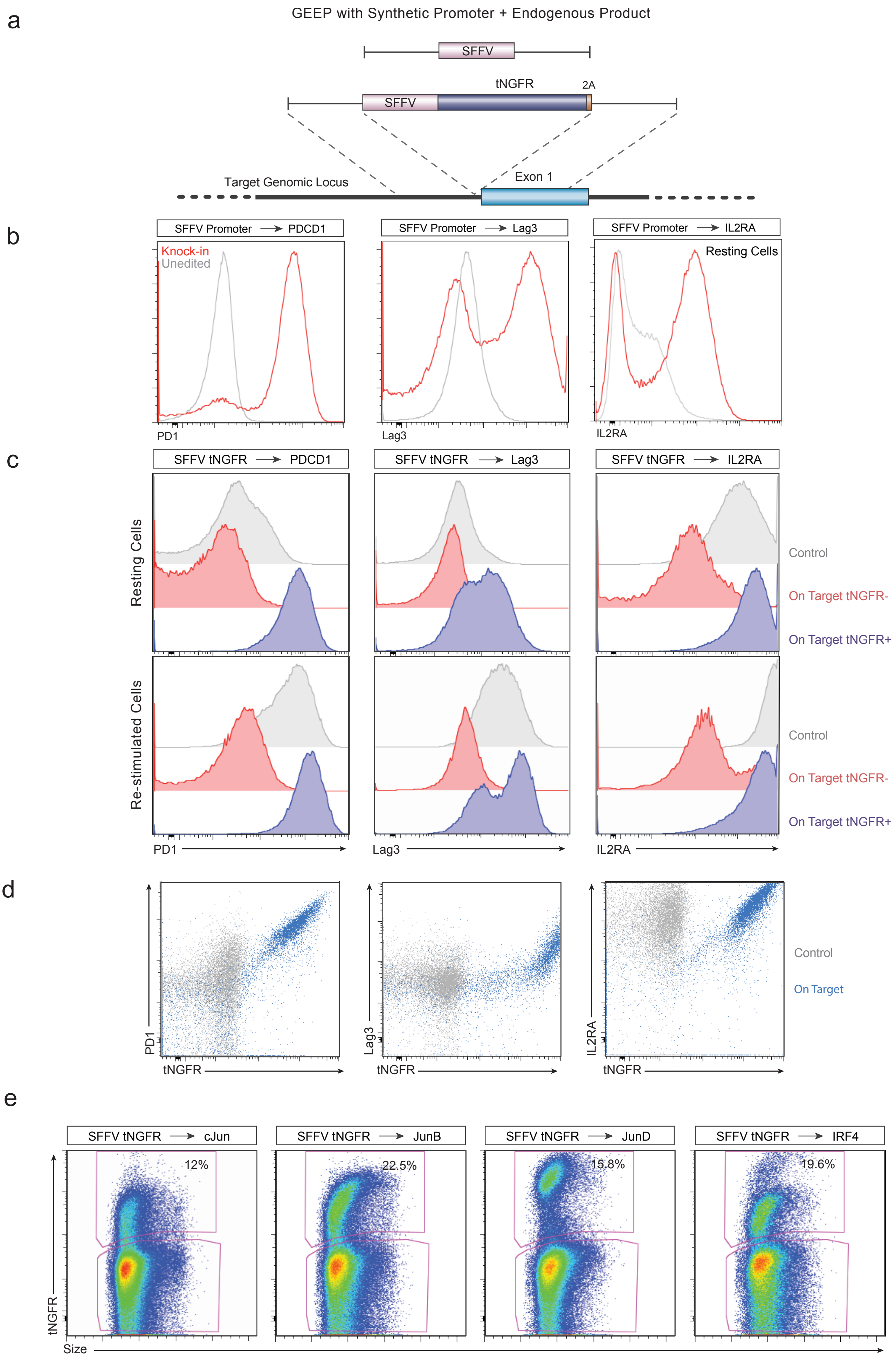


Extended Data Fig. 5

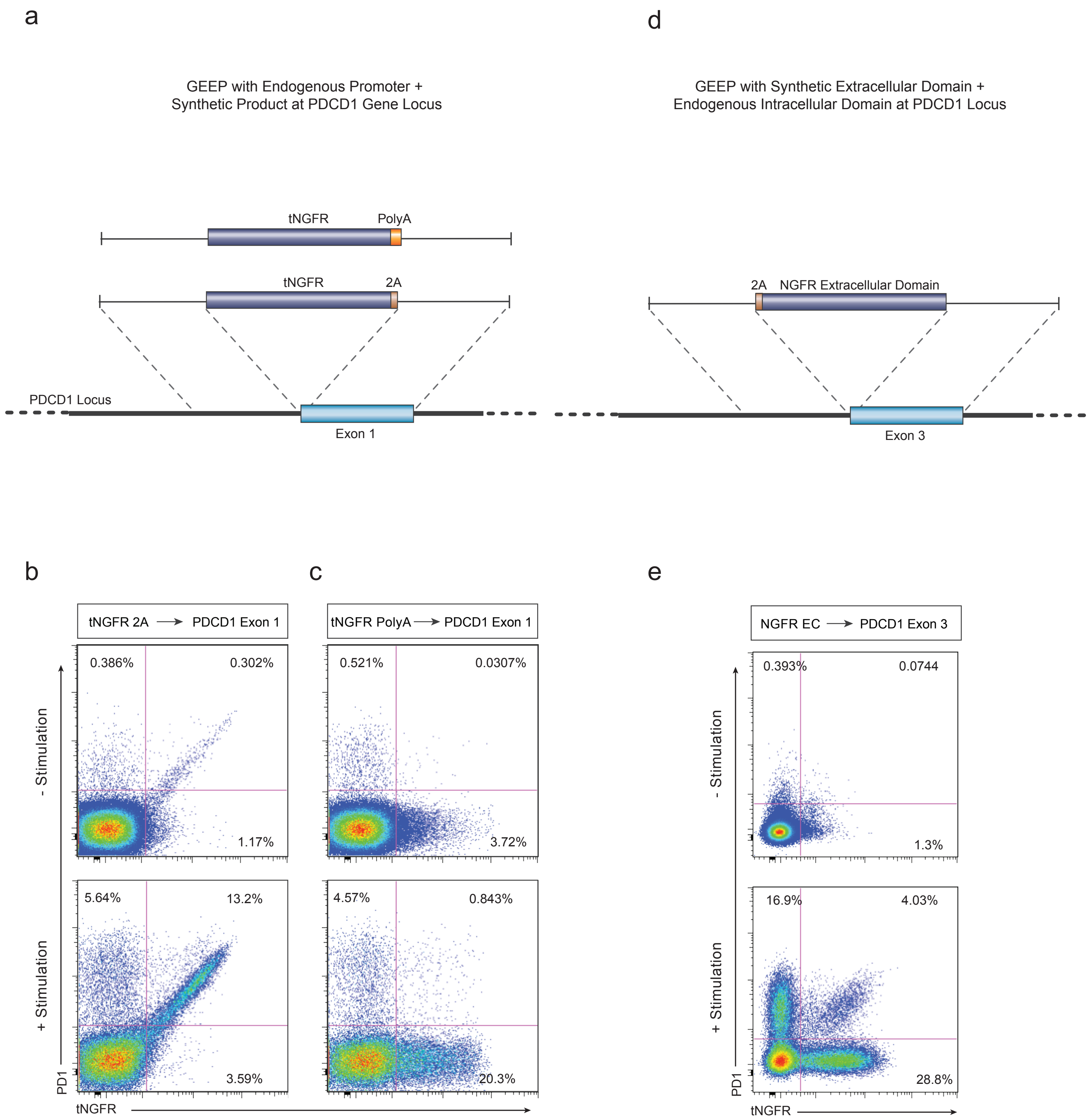


Extended Data Fig. 6

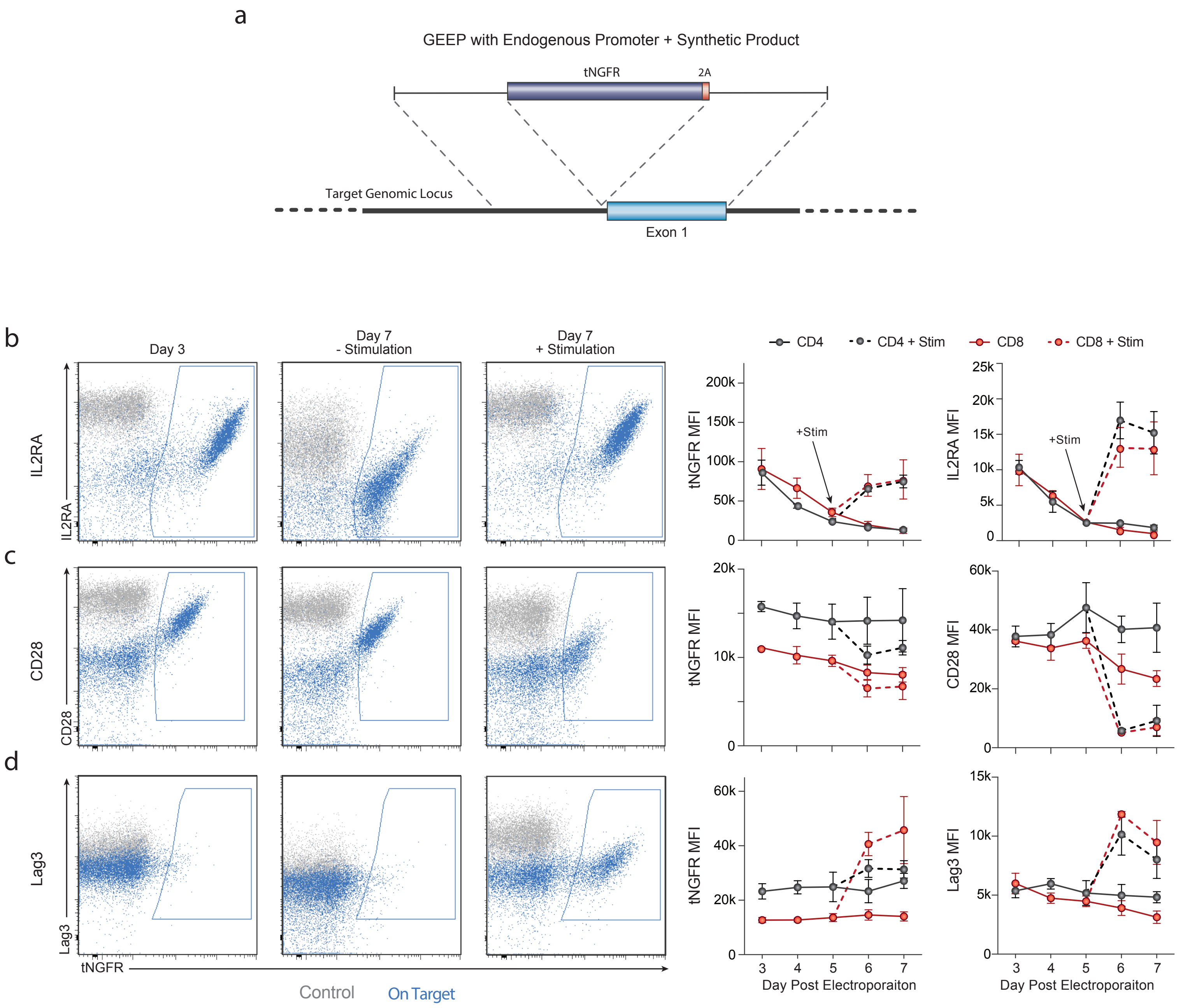


Extended Data Fig. 7

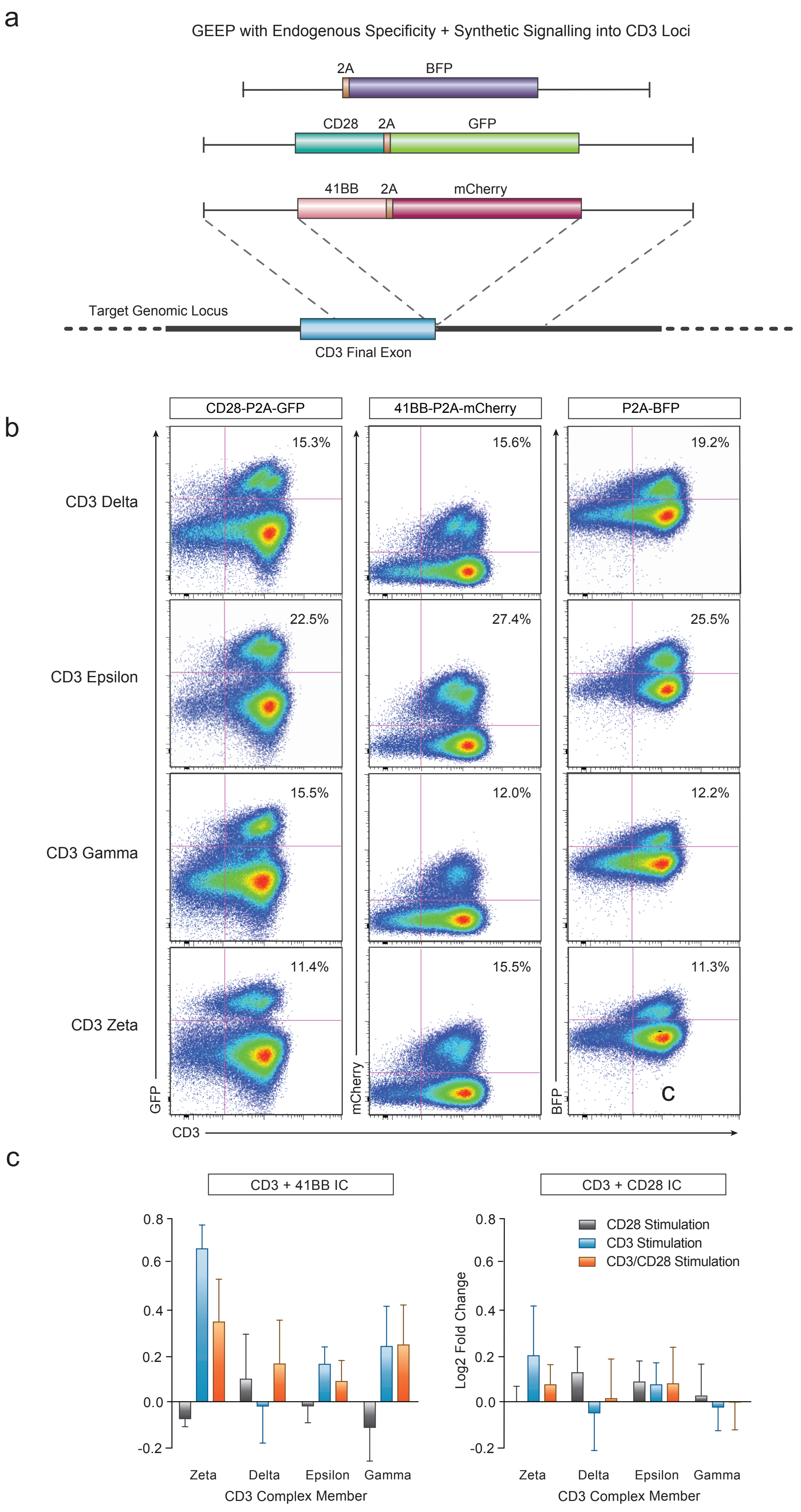


Extended Data Fig. 8

a

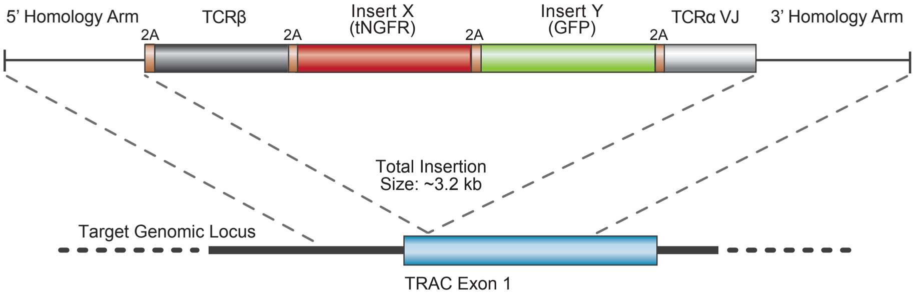


b

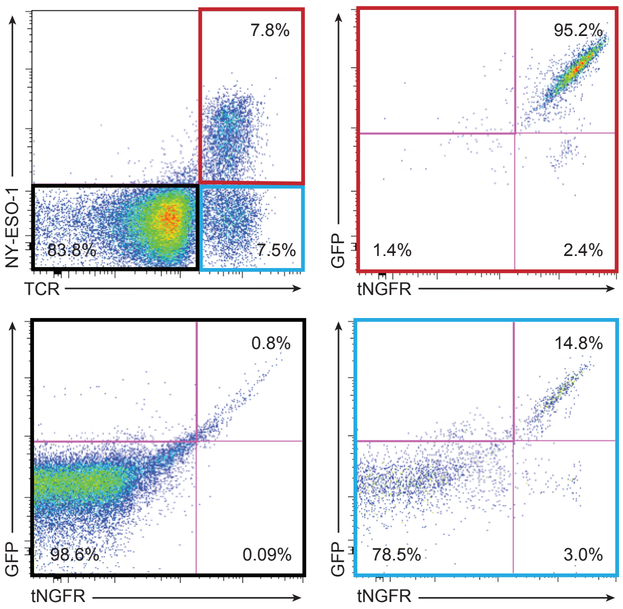


Extended Data Fig. 9

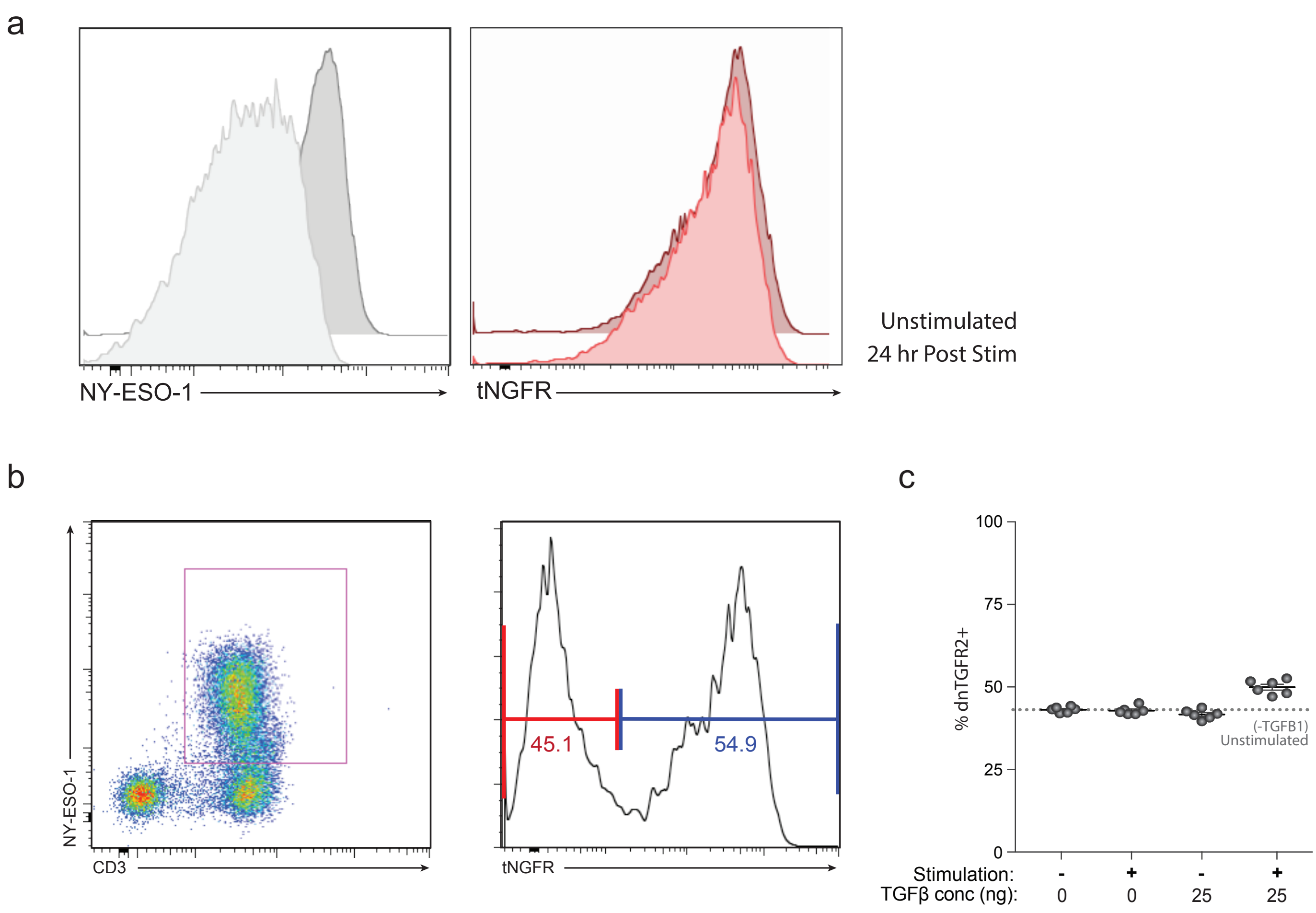


Extended Data Fig. 10

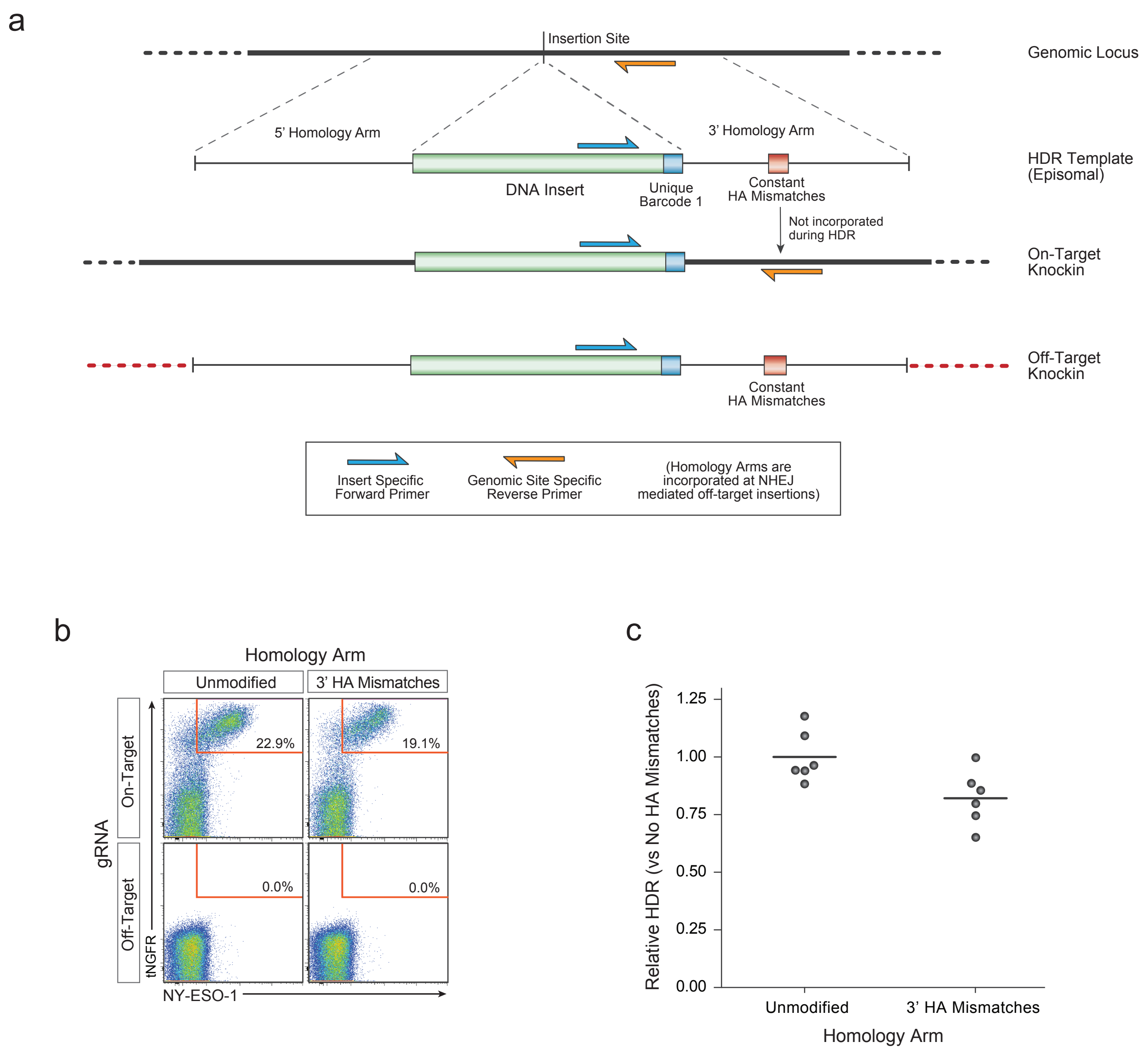


Extended Data Fig. 11

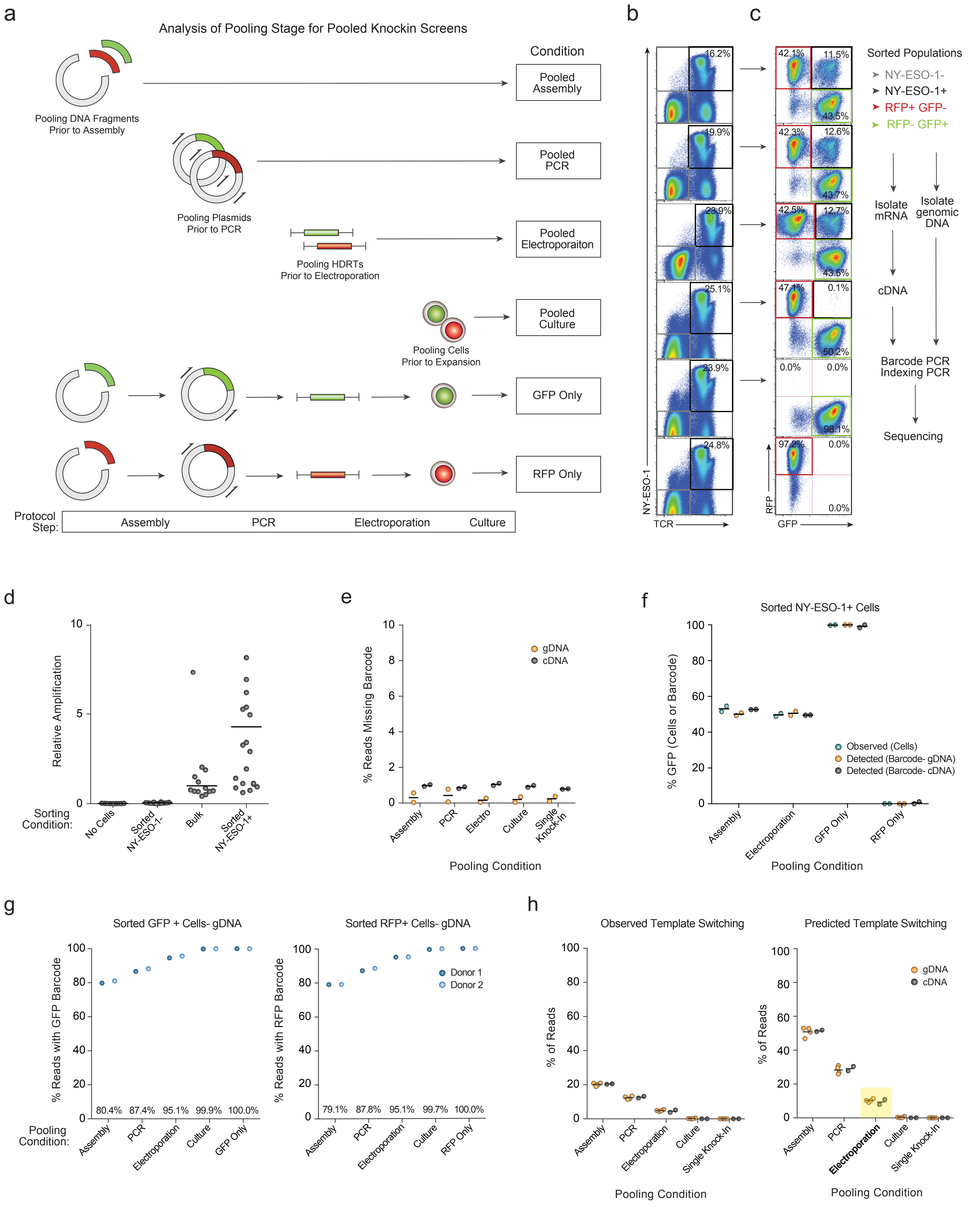


Extended Data Fig. 12

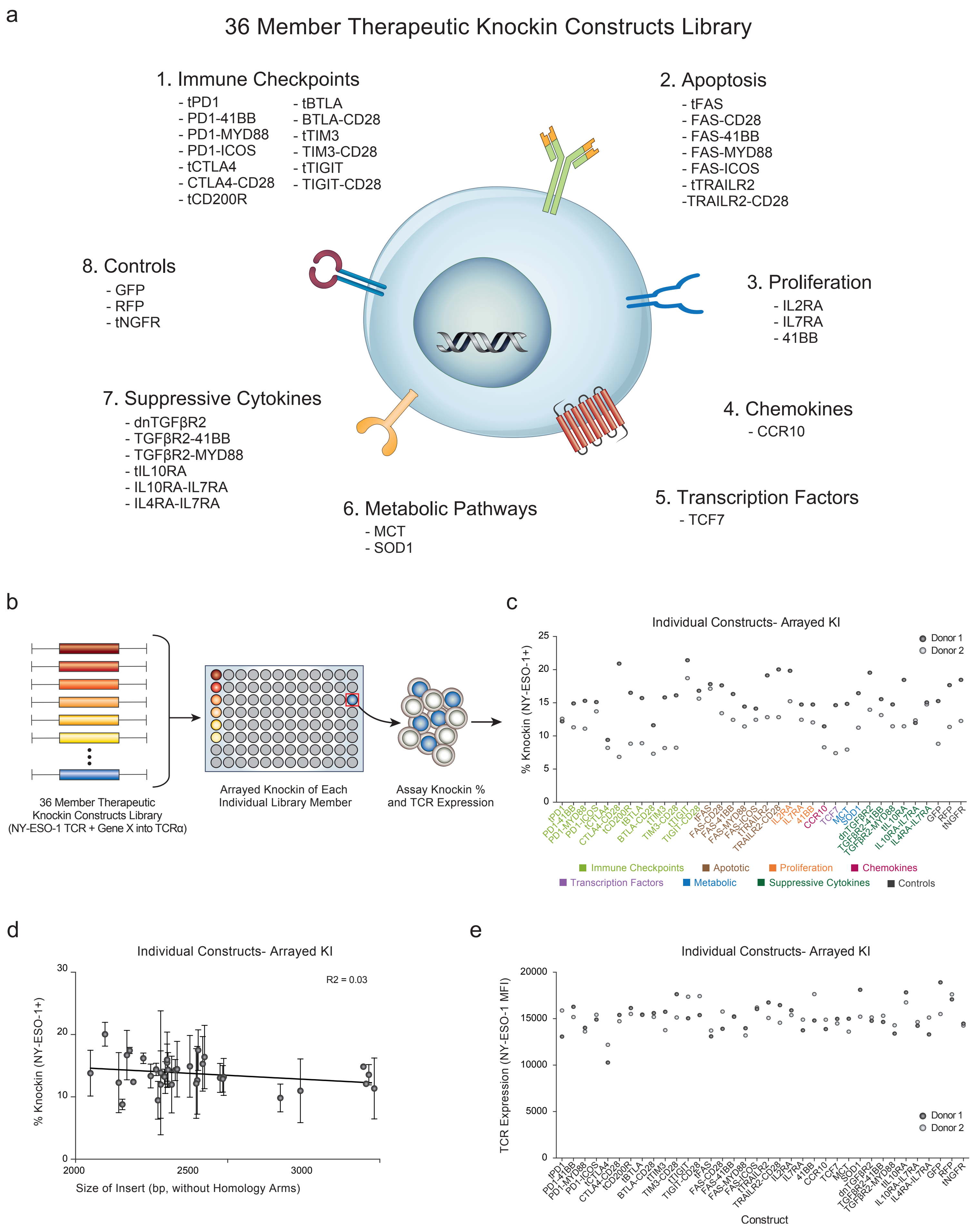


Extended Data Fig. 13

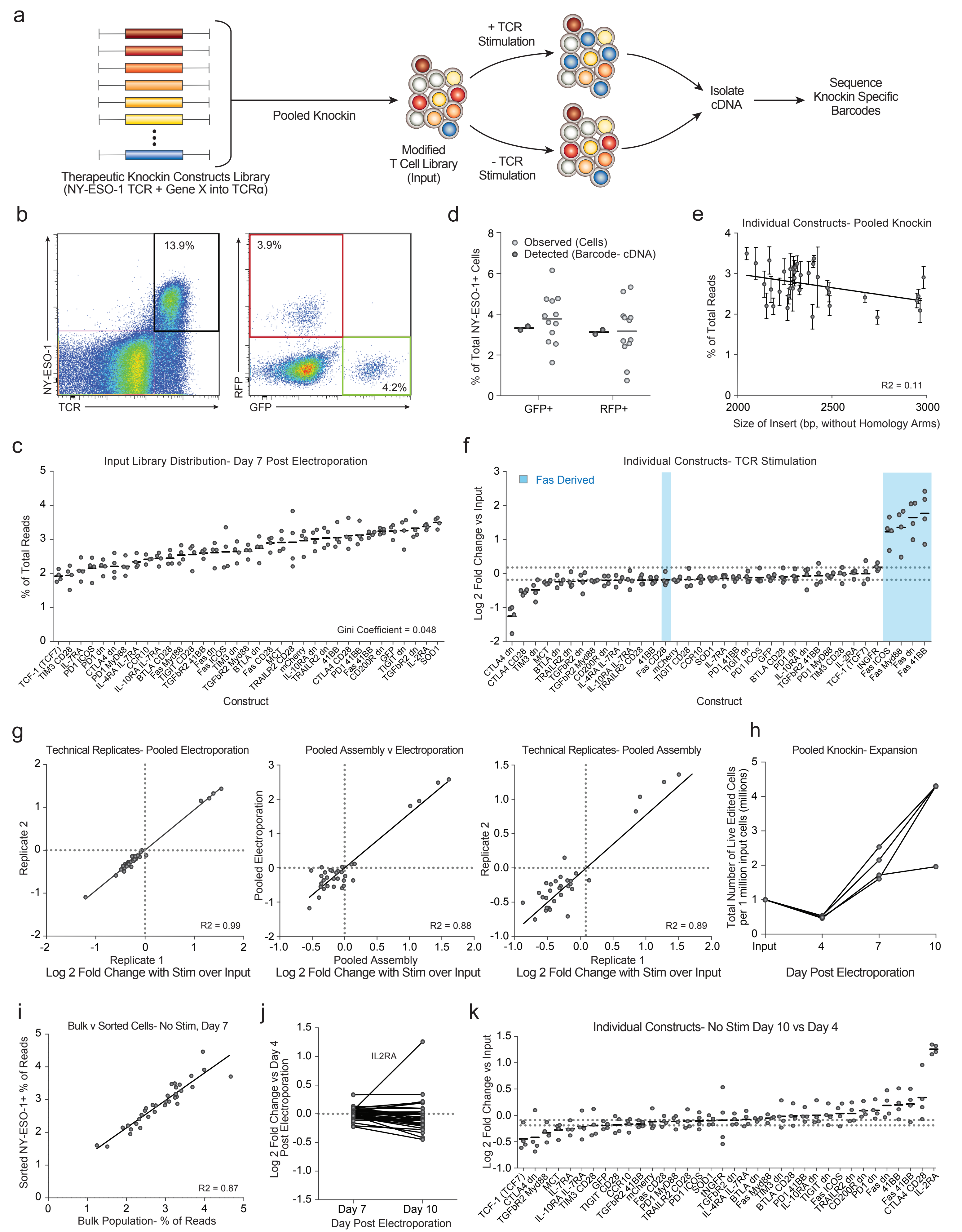


Extended Data Fig. 14

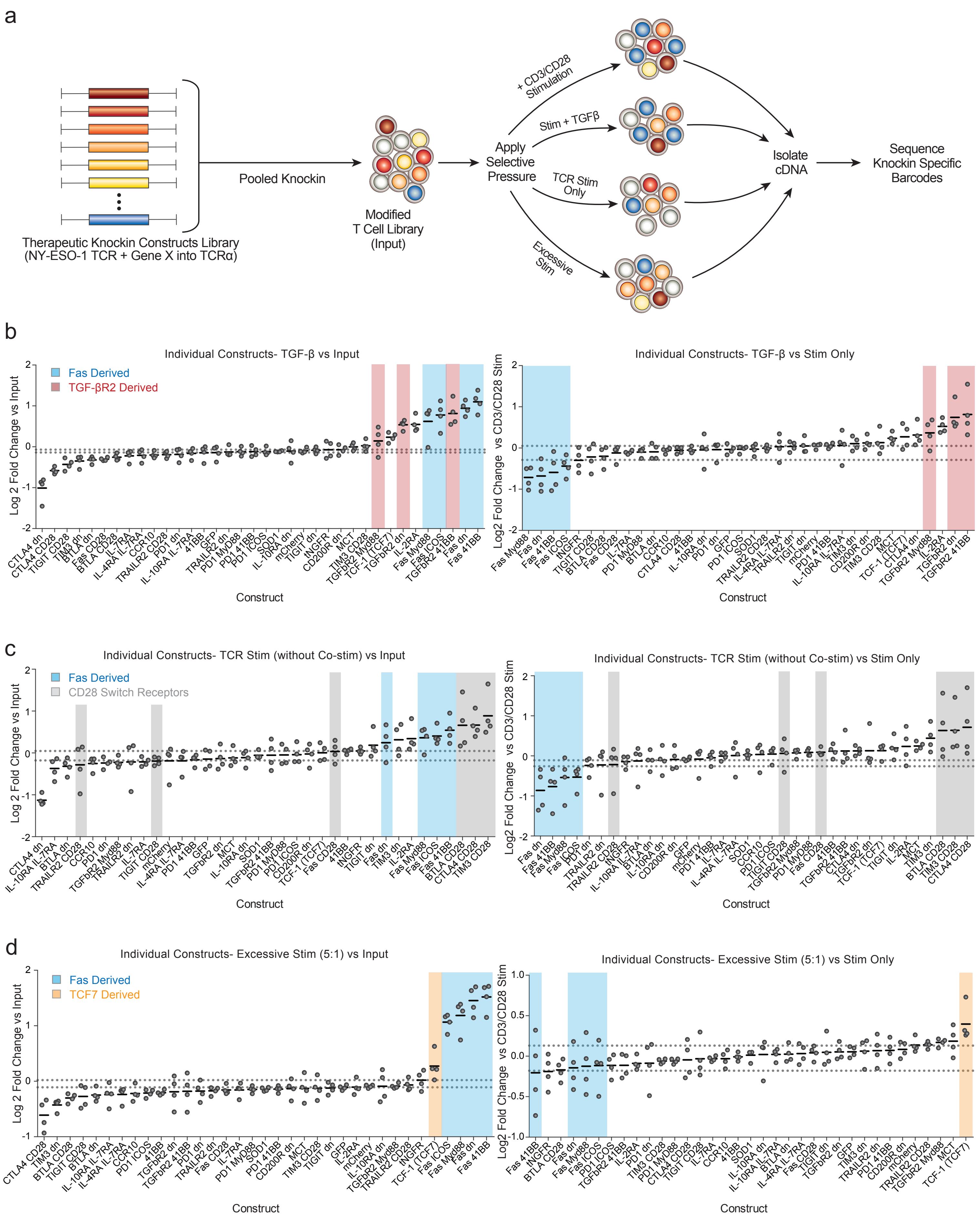


Extended Data Fig. 15

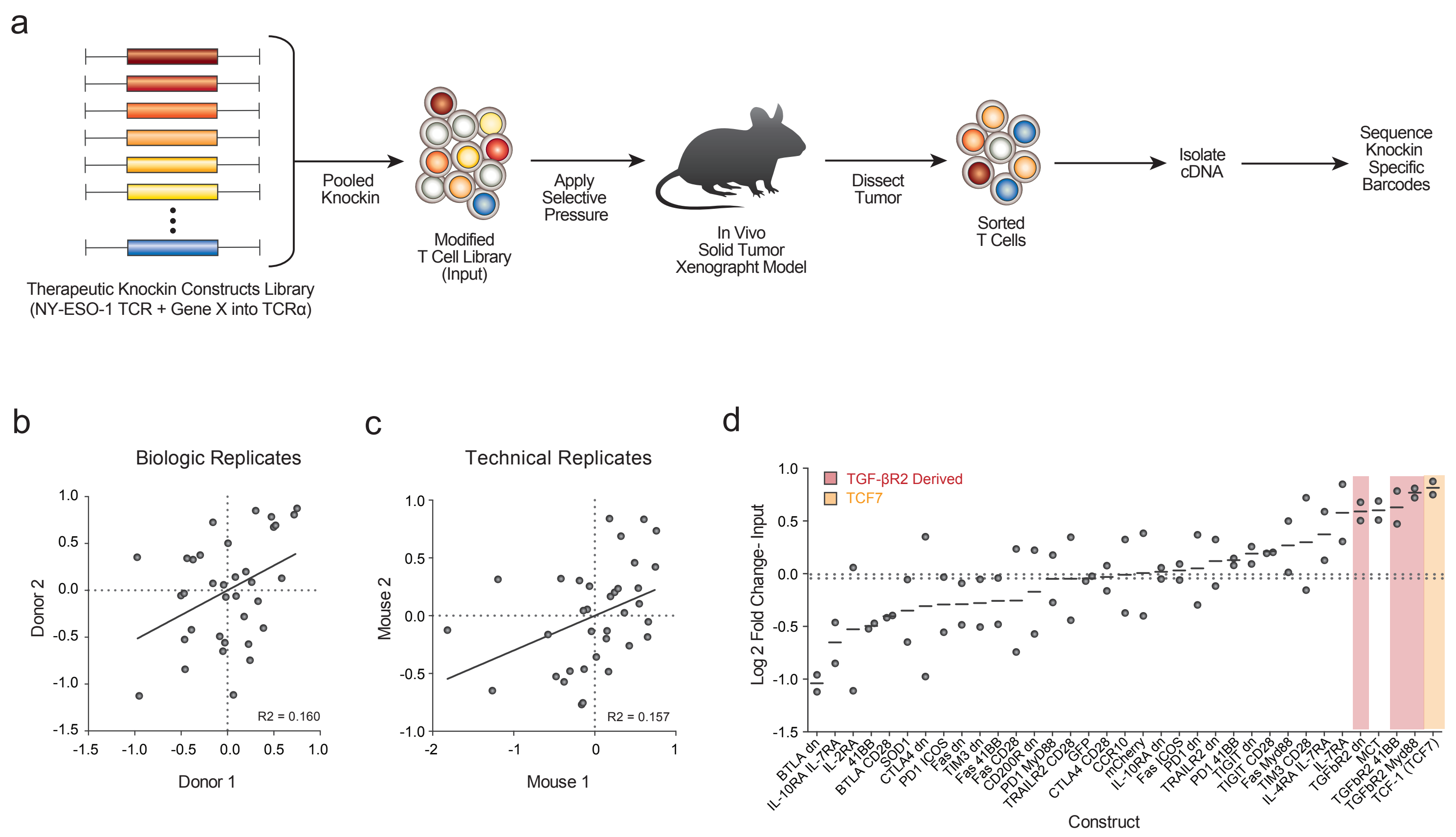


Extended Data Fig. 16

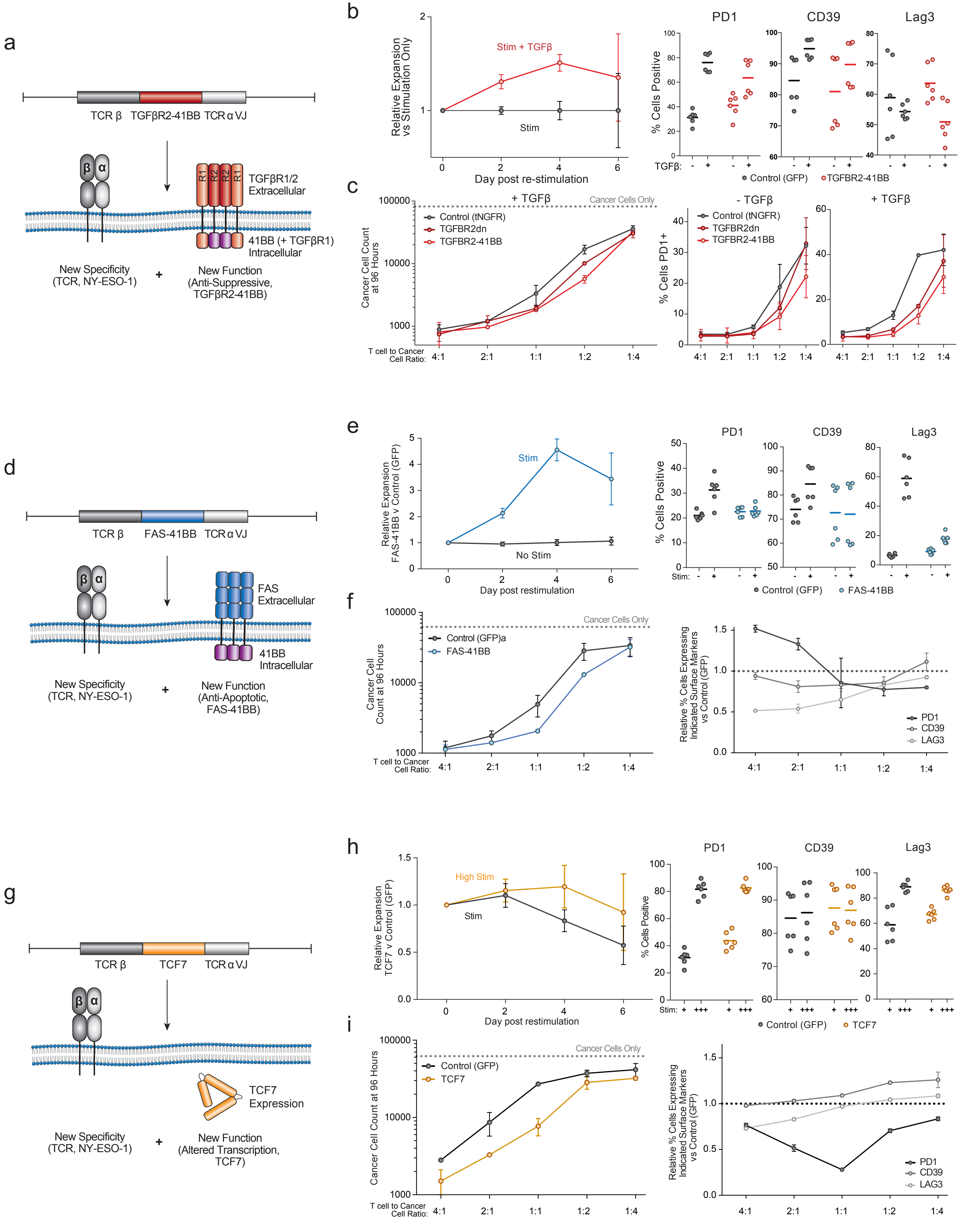


Extended Data Fig. 17
